# Beyond Antagonism: IL-4 Exploits TNF signaling to Shape Its Gene Expression Signature in Monocytes and Macrophages

**DOI:** 10.1101/2025.07.28.667180

**Authors:** Ruoxi Yuan, Chao Yang, Bikash Mishra, David Oliver, Richard Bell, Lionel B. Ivashkiv

**Affiliations:** HSS Research Institute and David Z. Rosensweig Genomics Research Center, Hospital for Special Surgery, New York, NY; Immunology and Microbial Pathogenesis Program, Weill Cornell Medicine, New York, NY; Department of Medicine, Weill Cornell Medicine, New York, NY

## Abstract

Investigation of crosstalk between antagonistic pro- and anti-inflammatory cytokines has focused on mechanisms and functional consequences of cross-inhibition. We investigated cross-regulation between proinflammatory TNF and anti-inflammatory IL-4 in primary human monocytes and in a skin wound-healing model. Surprisingly, TNF functioned mainly as a costimulator of IL-4-induced gene expression, whereas IL-4 selectively inhibited the TNF-induced IFN response, leaving inflammatory gene expression mostly intact. TNF and IL-4 synergistically induced gene sets important for regulating inflammation and tissue repair, which were highly induced during the phase of wound healing when these cytokines are co-expressed. Crosstalk between TNF and IL-4 was mediated by epigenetic chromatin-mediated mechanisms associated with cooperation between NF-κB and STAT6 transcription factors, erasure of negative histone mark H3K27me3, and selective inhibition of IRF1. These results identify a long-sought mechanism for expansion of the IL-4 response, and highlight the complexity of crosstalk between antagonistic cytokines that includes cooperation for select gene responses important in immune response and tissue repair.

## INTRODUCTION

Type 2 immune responses are characterized by the activity of the cytokines interleukin-4 (IL-4), IL-5, IL-9, and IL-13^1^. In addition to its classical function in host defense against parasitic and helminth pathogens, type 2 immunity serves various protective functions in the host, encompassing the regulation of metabolic equilibrium, suppression of excessive type 1 inflammation, fortification of barrier defenses, and the facilitation of tissue repair and regeneration ^2^. Notably, excessive inflammation hampers the tissue repair process, making the control of inflammation and transition from type 1 antimicrobial responses to type 2 tissue-reparative responses a critical step in restoring tissue homeostasis ^3^. IL-4 has anti-inflammatory effects, which it exerts primarily through signaling via the IL-4 receptor (IL4R), making it a pivotal contributor to tissue repair within the realm of type 2 immunity. As such, IL-4 plays an important role in skin wound healing, a complex, multi-stage process that begins immediately upon injury. It involves various skin cells like keratinocytes, fibroblasts, and endothelial cells, along with recruited immune cells ^4^. At the core of this intricate orchestration are monocytes and macrophages, which play central roles in directing the complex process of tissue repair ^5^. Their ability to undergo significant phenotypic, functional, and metabolic adaptations in response to local signals profoundly influences the surrounding cellular environment and the processes of tissue repair.

The IL-4 receptors consist of two types: type I IL-4R, comprised of IL-4Rα and γc subunits, and type II IL-4R, comprised of IL-4Rα and IL-13Rα1. IL-4 signals through both type I and type II IL-4 receptors, while its functionally closely related cytokine, IL-13, exclusively utilizes the type II receptor ^6^. Both receptors activate the JAK/STAT6 and PI3K signaling pathways in myeloid cells ^5^. Once activated, STAT6, the primary transcription factor associated with IL-4 signaling, plays a pivotal role in facilitating the transcriptional activation of genes including mannose receptor 1 (MRC1), CD23 (FCER2, FcR for IgE), arachidonate 15-lipoxygenase (ALOX15), as well as the chemokine genes C-C Motif Chemokine Ligand 17 (CCL17) and CCL22 ^7^. Although alternative signaling pathways have been identified, it’s noteworthy that STAT6-deficient mice typically phenocopy the effects of IL-4Rα deficiency ^8, 9, 10^. In addition to direct STAT6-mediated transcriptional activation of immune effector genes, IL-4 induces expression of a transcription factor cascade that further expands the IL-4-mediated gene expression program; key transcription factors activated downstream of IL-4 include IRF4, PPARγ, and EGR2 ^11, 12, 13^. An important associated mechanism of IL-4 action is epigenomic regulation, which encompasses DNA methylation, histone modifications, and chromatin remodeling ^14^. IL-4 utilizes these transcriptional and epigenetic mechanisms to fine-tune cellular responses to IL-4 itself and to modulate responses of cells to distinct signaling pathways. For instance, IL-4 has been shown to attenuate parts of the interferon-gamma (IFN-γ) response by targeting the auxiliary transcription factors, such as AP-1, that cooperate with the core IFN-γ-induced STAT1 and IRF1 factors ^15^. Moreover, STAT6 exerts repressive effects on a subset of enhancers that display a significant overlap with the NF-κB p65 cistrome ^16^. This interaction results in reduced responsiveness to lipopolysaccharide (LPS) following IL-4 stimulation. However, modulation of the TLR4 response by IL-4 is complex, with dichotomous regulation of various genes regulated in large part by transcription factor EGR2 ^11^. In contrast to regulation of distinct signaling pathways by IL-4, little is known about how IL-4 signaling itself is regulated by environmental signals. Notably, IFN-γ attenuates parts of the IL-4 response via binding of STAT1 to regulatory elements associated with IL-4-inducible genes ^15^. Additionally, granulocyte-macrophage colony-stimulating factor (GM-CSF) contributes to the IL-4-mediated up-regulation of IRF4 and its downstream target, CCL17 ^17, 18, 19^. Two landmark studies showed that IL-4 alone, *in vitro* and *in vivo*, elicits a muted response and requires co-stimulation for robust gene induction that is important for the function of IL-4 in type 2 responses, lung homeostasis, helminth and Listeria infections, and colitis ^20, 21^. Costimulation of IL-4 responses was provided in a context-specific manner by surfactant protein A, C1q, or apoptotic cells, which enhanced expression of various anti-inflammatory and wound-healing genes. Despite the importance of such an amplification system for IL-4 responses, little is known about underlying mechanisms.

IL-4 is co-expressed with the potent pro-inflammatory cytokine TNF in various biological contexts, including during the early stages of skin wound healing ^5, 22, 23, 24, 25^. In this study, we sought to test whether TNF modulates the IL-4 response, anticipating that TNF would suppress a subset of IL-4 induced genes, similar to suppression mediated by IFN-γ. Surprisingly, we found instead that TNF functions as a co-stimulator of most IL-4-induced genes including genes important for regulating inflammation and tissue repair. Using an epigenomic analysis in primary human monocytes that are directly relevant for inflammatory processes, we identified synergy between transcription factors STAT6 and NF-κB p65 subunit as a mechanism for co-stimulation of IL-4 responses. IL-4 reciprocally regulated TNF signaling by selectively inhibiting IRF1 while leaving NF-κB signaling mostly intact. This suggests a mechanism by which IL-4 can alleviate inflammation by suppressing TNF-induced IFN responses, which are mediated by IRF1 ^26, 27, 28, 29^. Moreover, co-stimulation with TNF and IL-4 also induced *KDM6B*, leading to the removal of the repressive histone mark H3K27me3, thereby further contributing to the induction of synergy genes. Our study paves the way for a deeper understanding of epigenetic mechanisms that underlie the complex interplay between cytokines with simultaneous synergistic and antagonistic interactions, which holds promise for developing strategies to modulate inflammation and tissue repair.

## RESULTS

### Co-induction of TNF-NF-κB and IL-4 pathways in a subset of macrophages during skin wound healing

Skin wound healing progresses from an early inflammatory phase driven by cytokines such as TNF through proliferative and resolution phases when IL-4 is active ^5, 22, 23, 24, 25^. We identified an MHC class II-hi CD206-lo macrophage population in mouse skin wounds 4 dpi that expressed very high levels of IL-4-inducible genes *Ccl17*, *Ccl22*, *Irf4* and *Kdm6b* relative to MHC II-lo CD206-hi macrophages (Fig. 1a-1c; the gating strategy is shown in Extended Data Fig. 1a). These cells also showed higher levels of *Il1b* and of IL-4 target gene *Retnla* expression, whereas CD206-hi MHC II-lo macrophages expressed higher levels of *Arg1*, which is induced by IL-4 and other pathways ^11, 30^. Macrophages in skin wounds could also be subdivided into F4/80-high and F4/80-low subsets, with MHC II-hi macrophages expressing lower levels of F4/80 (Fig. 1d, e). F4/80-low macrophages expressed high levels of *Ccl17*, *Ccl22*, *Irf4*, *Alox15* and also *Il1b*, but low levels of *Cd206* relative to F4/80-high macrophages (Fig. 1f). These results show co-expression of select IL-4 target genes and inflammatory NF-κB target *Il1b* in an MHC II-hi CD206-low F4/80-low macrophage subset *in vivo* during skin wound healing.

**Fig. 1.**
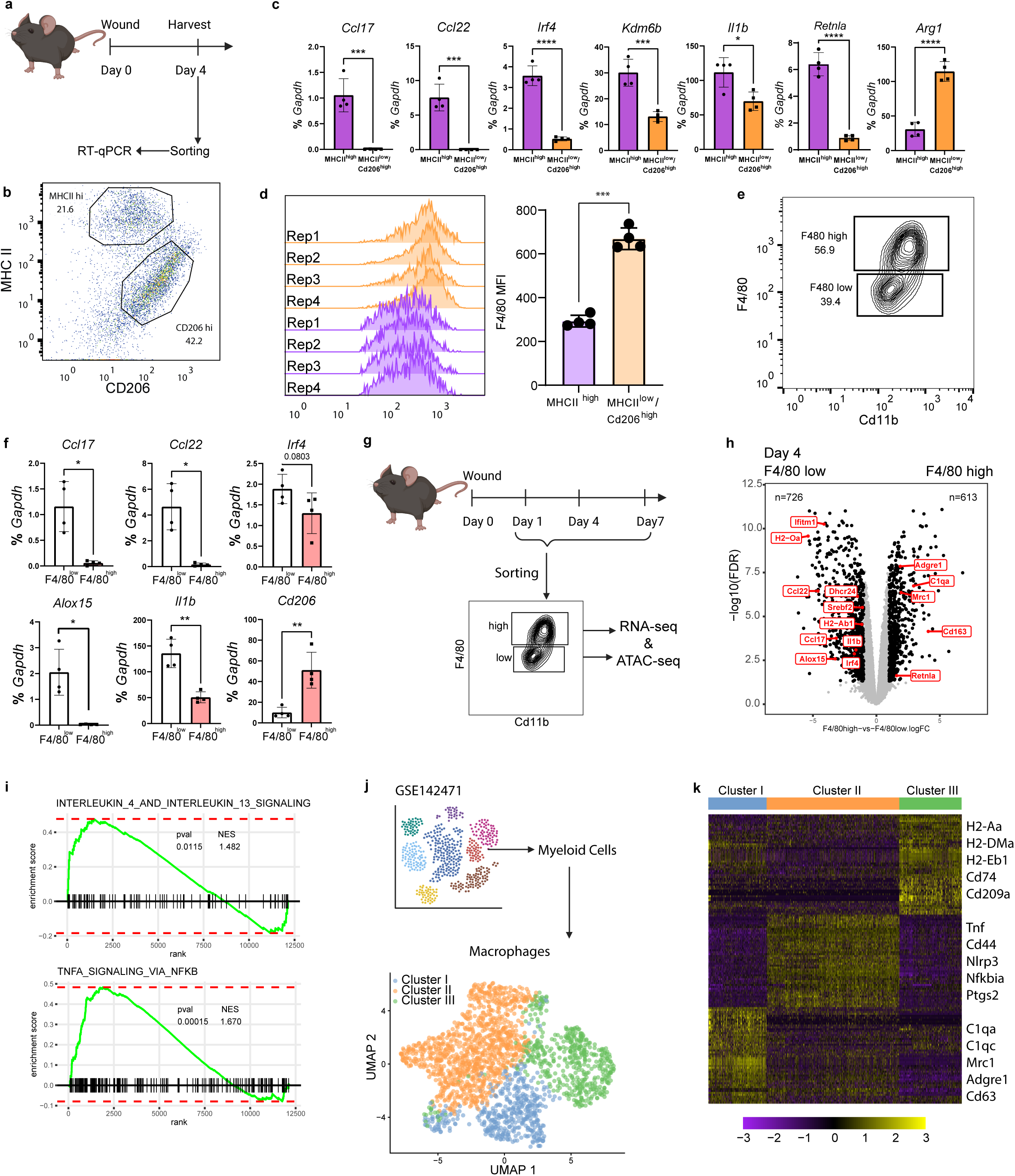
Distinct macrophage phenotypes in mouse skin during wound healing. **a**, Schematic illustration of the experimental design. **b**, Representative flow cytometry plot of 4 independent experiments for skin wound macrophage subsets defined by MHCII and CD206 expression, also see gating strategy in Extended Data Fig 1a. **c**, RT-qPCR analysis for indicated genes normalized relative to *Gapdh* of flow-sorted MHCII high and MHCII low/CD206 high cells, as depicted in Fig 1a, at 4 dpi, n = 4. Each dot represents a sample of pooled cells from 3 mice per experiment. Representative of 2 independent experiments. *p value < 0.05, ***p value < 0.001, ****p value < 0.0001, paired t test. Data are presented as means ± SD. **d**, Left. Flow cytometry histogram plot of F4/80 cell surface expression in CD206 hi and MHCII hi monocytes at day 4 post injury. 4 independent replicates are depicted. ***p value < 0.001, paired t test. Right. Data are presented as means ± SD. **e**, Flow cytometry plot of F4/80 high and F4/80 low macrophages, representative of 4 independent experiments, also see gating strategy in Extended Data Fig 1a. **f**, RT-qPCR analysis of indicated gene expression normalized to *Gapdh* of sorted F4/80 high and F4/80 low cells as indicated in Fig 1e, at 4 dpi. *p < 0.05, **p < 0.01, ns: not significant. Paired t test was used. Data are presented as means ± SD. **g**, Schematic illustration of the experimental design for RNA-seq and ATAC-seq. **h**, Volcano plots display the statistical significance (-log10 FDR, y-axis) versus the log2 fold change of RNA-seq data, highlighting selected genes that were more highly expressed in F4/80-high macrophages (right) and F4/80-low macrophages (left) at 4 dpi. **i**, GSEA results of ranked differentially expressed genes from RNA-seq data in Fig. 1h (n = 3) using the REACTOME_INTERLEUKIN_4_AND_INTERLEUKIN_13_SIGNALING and HALLMARK_TNFA_SIGNALING_VIA_NFKB gene set. The Normalized Enrichment Score (NES) provide a normalized measure of the degree to which a set of genes is overrepresented at the top or bottom of a ranked list of genes. Values of p value < 0.05 were considered statistically significant. **j**, UMAP projection of macrophage subsets from scRNA-seq dataset of skin wounds at 4 dpi from GSE142471. Cluster I: MHCII^lo^ Cd206^hi^ F4/80^hi^ population; Cluster II: MHCII^lo^ Cd206^lo^ F4/80^lo^ population; Cluster III: MHCII^hi^ Cd206^lo^ F4/80^lo^ population. **k**, Heatmap depicting the top 50 differentially expressed genes in each cluster. Columns represent single cells, with differential gene expression scaled from low (purple) to high (yellow).

To gain greater insight into co-activation of IL-4 and TNF-NF-κB pathways in skin wound macrophages, we flow-sorted F4/80-high and F4/80-low macrophages at day 1, day 4, and day 7 post-injury and performed RNAseq and ATACseq (Fig. 1g, Extended Data Fig. 1b). Principal component analysis of RNAseq data (Extended Data, Fig. 1b) showed that day 1 macrophages segregated from day 4 and 7 macrophages, and that on days 4 and 7 macrophages segregated primarily based on F4/80-low versus F4/80-high phenotype. In line with these results, hierarchical clustering of differentially expressed genes (DEGs) revealed a gene cluster (cluster 1) that was most highly induced in day 1 F4/80 low macrophages and contained *1l1b* (Extended Data, Fig. 1c). Cluster 1 showed highly significant enrichment of genes in NF-κB, inflammatory response, hypoxia and IFN response pathways (Extended Data, Fig. 1d), consistent with a predominant type 1 response at this time point. Expression of these genes was partially sustained at day 4 and to a lesser extent on day 7. In contrast, expression of cluster 2 genes peaked at day 4 in F4/80-low macrophages and was sustained on day 7. Cluster 2 showed enrichment of allograft rejection and cholesterol homeostasis genes (Extended Data Fig. 1d) and notably contained the above-described subset of IL-4-inducible genes such as *Alox15*, *Ccl17* and *Ccl22*, and MHC genes (Extended Data Fig. 1c and Fig. 1h). Regulation of cholesterol pathway genes such as *Dhcr24* and *Srebf2* is consistent with a late phase TNF response ^26^. Strikingly, gene set enrichment analysis (GSEA) of the F4/80-low population revealed significant enrichment in pathways related to both IL-4 signaling and TNF-mediated NF-κB signaling (Fig. 1i). A decrease in MHC II-high and F4/80-low cells in TNF knockout mice or after inhibition of STAT6 (Extended Data Fig. 1e, 1f) further supported a role for TNF and IL-4 signaling in induction of this cell phenotype. In contrast to F4/80-low cells, F4/80-high macrophages expressed a distinct set of IL-4-inducible genes such as *Retnla*, *Cd36*, *Pparg*, and *Mrc1*, were enriched for genes involved in collagen remodeling ^31^ and cell proliferation, and showed decreased expression of inflammatory pathways (Extended Data Fig. 1c, 1d, 1g-1i). These cells expressed canonical markers of tissue-resident macrophages, such as *C1qa*, *Apoe*, *Folr2* and *Mertk*, in contrast to F4/80-low cells that expressed monocyte genes such as *Ly6c2*, *Ccr2* and *S100a9*. These results suggest that TNF and IL-4 pathways converge in F4/80-low macrophages at 4 days after skin wounding. These F4/80-low macrophages express inflammatory, MHC class II and a subset of IL-4 target genes, but the role of TNF and IL-4 pathways in regulating these genes, and whether these cytokines antagonize each other, as predicted, is not clear.

To further corroborate these findings, we re-analyzed a previously reported dataset ^32^ of single-cell RNA sequencing (scRNA-seq) data from cells isolated from skin wounds at 4 days post-injury (dpi) (Extended Data Fig. 2a, 2b). We subsetted cells expressing *Ptprc* (CD45) from the original dataset and further clustered them into four distinct populations based on marker gene expression. Subclustering of macrophages revealed three broad clusters (Fig. 1j). Strikingly, Cluster III macrophages that exhibited an MHC II-hi, CD206-lo F4/80-lo phenotype expressed enhanced IL-4-inducible genes (Fig. 1k and Extended Data, Fig. 2c).

We further examined ATAC-seq data from F4/80-high and F4/80-low macrophages to assess differences in chromatin landscapes (Extended Data Fig. 2d). Surprisingly, we observed only a limited number of significantly different ATAC-seq peaks between the two macrophage populations (Extended Data Fig. 2e). To gain insight into transcription factors that may regulate gene expression, we performed motif enrichment analysis on ATAC-seq peaks identified as enhancer regions for genes in cluster h2 that contained IL-4-inducible, MHC and inflammatory genes (Extended Data Fig. 1c). *De novo* motif enrichment results suggested potential involvement of STAT and NF-κB signaling pathways in regulating these genes (Extended Data Fig. 2f). To determine whether ATAC-seq peaks in F4/80-high and F4/80-low macrophages could be differentiated based upon transcription factor occupancy, we performed footprinting analysis on ATAC-seq data using TOBIAS. Interestingly, despite minimal significant differences in chromatin openness, the results indicated significantly increased binding of pro-inflammatory mediators such as REL and NFKB1 in F4/80-low macrophages at 1 dpi (Extended data Fig. 2g). Additionally, we observed enhanced binding signals of both NFKB1 and STAT6 in F4/80-low macrophages at 4 dpi, whereas the overall differences in transcription factor binding between F4/80-high and F4/80-low macrophages diminished by 7 dpi (Extended Data Fig. 2g). This result was further validated by ChromVAR at 4 dpi (Extended Fig. 2h). These results suggest that the overall skin wound environment imparts a similar chromatin accessibility landscape on both populations of macrophages. However, stronger and concomitant activation of NF-κB and IL-4 pathways was observed in F4/80-low macrophages that expressed NF-κB and IL-4 target genes, further supporting an important role for NF-κB-IL-4-STAT6 signaling crosstalk. To investigate functional consequences of NF-κB and IL-4 signaling crosstalk and underlying mechanisms, we transitioned to an *in vitro* system.

**Fig. 2.**
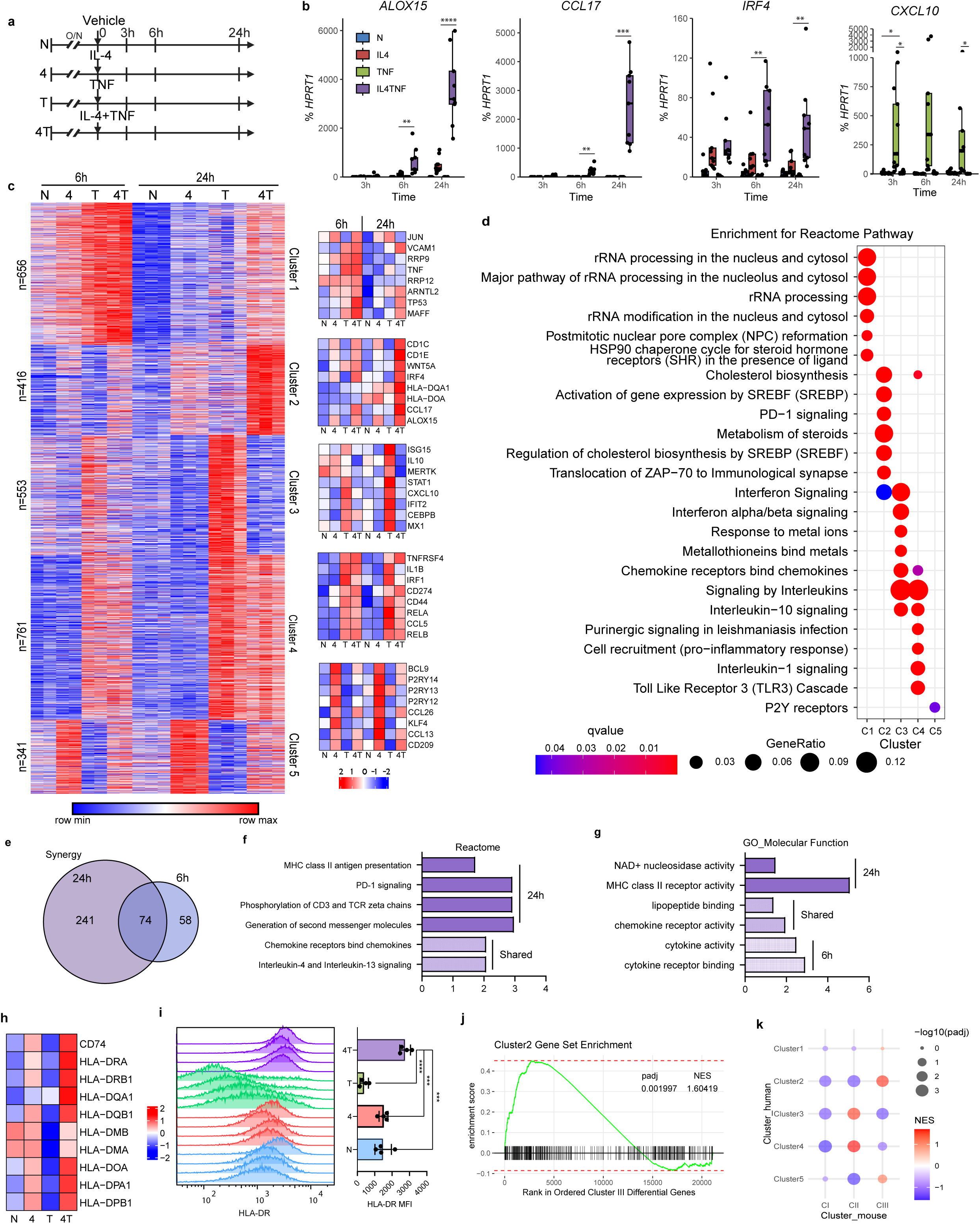
Synergistic and Antagonistic Interplay between IL-4 and TNF in Regulating Gene Expression in Human Monocytes. **a,** Experimental design: human CD14^+^ primary monocytes were stimulated with IL-4 (30ng/ml) and/or TNF (10ng/ml) for 3, 6 or 24 hours. **b**, RT-qPCR measurement of the indicated mRNA at the indicated time points after IL-4 and TNF costimulation (n=9). Boxplots displaying the interquartile range of expression levels for various treatments (N, IL4, TNF, IL4TNF) at different time points (3h, 6h, 24h). Each dot represents an individual donor, and the horizontal line within each box shows the median value. Two-way ANOVA was performed at each time point. *p < 0.05, **p < 0.01, ***p < 0.001. **c**, Left panel, K-means (K=5) clustering of 2727 differentially up-regulated genes (log2 fold change > 1 and FDR < 0.05) in any pairwise comparison to untreated control. Values from biological replicates are shown in separate columns for each sample (6h: n=2, 24h: n=3). Right panels show heatmaps of expression of representative genes in each cluster. **d**, Reactome pathway enrichment analysis of the gene clusters in Fig 2c. **e**, Venn diagram of the overlap between genes synergistically induced by IL-4 and TNF co-stimulation at 6 hours and 24 hours. **f**, **g**, Functionally enriched Gene Ontology (GO) categories Reactome pathways (**f**) and Molecular Function (MF) (**g**) and of the synergy genes in Fig. 2e. **h**, Heatmap of gene expression from same RNA-seq analysis as in Figure 1 displaying representative MHC class II genes, color-coded by z-scores. **i**, Left: Representative histogram showing Median Fluorescence Intensity (MFI) of HLA-DR on human CD14^+^ monocytes treated with indicated experimental conditions for 24 hours. Cells were gated on live singlets (using a FSC-W versus FSC-A gate). Right: Cumulative data (n = 4) shown as means ± SD. ***p < 0.001, ****p < 0.0001, One-way ANOVA. **j**, **k**. Gene set enrichment analysis (gene Clusters from Figure 2c). Mouse wound healing clusters: CI: Cluster I, CII: Cluster II, CIII: Cluster III.

### Amplification of IL-4-induced gene responses by TNF

As the cells that exhibited co-activation of TNF-NF-κB and IL-4 pathways were most likely monocyte-derived, we wished to investigate mechanisms of TNF-IL-4 crosstalk in monocytes. We hypothesized that TNF would mostly antagonize IL-4 responses and thus attenuate IL-4-induced gene expression. Given limitations on mouse monocyte cell numbers, we assessed the effects of TNF on IL-4-induced gene expression in a time-course experiment using primary human monocytes (Fig. 2a). Surprisingly, TNF co-treatment strongly superinduced IL-4-activation of *ALOX15*, *CCL17*, and *IRF4* in a manner that increased over time and was most apparent at the 24 hr time point (Fig. 2b), suggesting TNF may actually contribute to the gene expression profile observed in F4/80-low skin wound macrophages. In contrast, IL-4 treatment strikingly suppressed *CXCL10* expression induced by TNF (Fig. 2b). Super-induction of IL-4 target genes was relatively specific for TNF relative to other inflammatory factors tested (Extended Data Fig. 3a) and was also observed when cells were preincubated with TNF or IL-4 before addition of the second cytokine (Extended Data Fig. 3b, c). To gain broader insights into gene regulation we performed RNA-seq at the 6 and 24 hr time points (Extended Data Fig. 3d, e). Analysis of differentially expressed genes (DEGs) (log fold change > 1, FDR < 0.05) based on pattern of expression revealed 5 distinct clusters (Fig. 2c). In contrast to expectations, expression of a majority of IL-4-inducible genes was further increased by TNF co-stimulation (Fig. 2c, clusters 1 and 2). Expression of cluster 1 genes was also increased by TNF alone and peaked at 6 hr, while expression of cluster 2 genes was minimally affected by TNF alone and peaked at 24 hr. To gain insight into the biological relevance of cooperation between TNF and IL-4, we performed pathway analysis and found that cluster 1 was enriched in ribosomal RNA processing genes, while cluster 2 was enriched in genes involved in cholesterol, fatty acid and steroid metabolism, which have been linked to regulation of inflammatory and interferon (IFN) responses ^26, 33, 34, 35^, in immune cell and PD-1 signaling, and in MHC class II antigen presentation (Fig. 2c, 2d and Extended Data Fig. 3f). The right panels of Fig. 2c display heatmaps of representative genes in each cluster. We also used a recently described strategy ^36^ to identify synergy genes whose induction by co-stimulation by both stimuli is at least 1.2-fold greater than the additive effects of individual stimuli (Fig. 2e, Extended Data Fig. 3g and Supplementary Table 1). When applying an FDR < 0.05, we found a total of 373 genes that were synergistically induced by TNF + IL-4 co-stimulation (Fig. 2e). Pathway analysis of this gene set, termed ‘synergy genes’ revealed enrichment in IL-4, chemokine, immune cell, and PD-1 signaling, and MHC class II antigen presentation pathways (Fig. 2f, g), mirroring the above-described analysis of cluster 2. We further identified a specific subset of “*de novo*” induced genes that were activated exclusively by co-stimulation, and not by individual cytokines (Supplementary Table 1). Interestingly, 37% of synergy genes were induced *de novo*, indicating that co-stimulation does not solely amplify, but also extends cytokine-induced gene responses, although pathway analysis showed that de novo genes reflected a similar biology as did synergy genes (Supplementary Table 1 and not shown). The majority of MHC class II genes were synergistically induced by IL-4 and TNF, which also suppressed CD206 expression (Fig. 2h, i and Extended Data Fig. 3i).

The induction of an HLA-DR-high, CD206-low phenotype in human monocytes, along with co-induction of select IL-4 target genes such as *ALOX15*, *CCL17* and *CCL22*, suggested similarity to the F4/80-low cell phenotype observed in mouse skin wounds at 4 dpi (Fig. 1f). To systematically assess whether the TNF + IL-4-induced genes identified in Fig. 2c were expressed in wound-healing macrophages, we performed GSEA. This analysis revealed highly significant enrichment of human Cluster 2 genes (Fig. 2c) within the wound-healing macrophage Cluster III identified in Fig 1j, 1k (Fig. 2j). In contrast, genes induced by TNF alone (Clusters 3 and 4, Fig. 2c) were specifically enriched in Cluster II (Fig. 2k). These findings highlight the cooperative role of TNF inflammatory pathways and IL-4 in activating monocyte and macrophage gene expression. Collectively, these results show that TNF broadly reprograms the transcriptional response to IL-4 in a mostly positive direction, increasing expression of genes in pathways related to immune responses, steroid metabolism and antigen presentation.

### Suppression of TNF-induced interferon response by IL-4

In contrast to predominantly positive regulation of IL-4-inducible genes by TNF, the majority of TNF-inducible genes were suppressed by IL-4 (Fig. 2c, clusters 3 and 4). Cluster 3 TNF-inducible genes that were strongly suppressed by IL-4 were enriched in interferon-stimulated genes (ISGs) (Fig. 2d and Extended Data Fig. 3f). This indicates that IL-4 suppresses the TNF-induced IFN response, which is mediated by an IRF1-dependent IFN-β autocrine loop ^26, 27, 29^. This attenuation of an IFN response mirrors suppression of the IFN response observed in skin would macrophages that occurred at the time of emergence of IL-4 signaling (Extended Data Fig. 1i), supporting a role in *in vivo* regulation. Cluster 4 TNF-inducible genes that were relatively weakly suppressed by IL-4 were enriched in genes related to IL-1 and NF-κB signaling (Fig. 2d). Additional Gene Set Enrichment Analysis (GSEA) confirmed that TNF expands and extends the IL-4 response, while IL-4 mainly suppresses the TNF-induced IFN response and also attenuates activity of AP-1/ATF transcription factors (Extended Data Fig. 3j). Overall, the results show that antagonism of the TNF response by IL-4 is strongly focused on the IFN autocrine loop, while TNF mostly augments and extends rather than inhibits the IL-4 response. Differences between IL-4 regulation of TNF- and TLR4-induced IFN responses may be related to the distinct mechanisms of IFN-β induction, mediated by, respectively, IRF1 and IRF3 ^26, 27, 29, 37, 38^.

### Regulation of chromatin accessibility and transcription factor activity

To gain insight into the mechanisms underlying TNF-IL-4 crosstalk, we conducted ATAC-seq to evaluate chromatin accessibility and analysis using ChromVAR to assess signal enrichment and transcription factor activity in open chromatin regions at 6 hours and 24 hours after cytokine stimulation (Extended Data Fig. 4a). Stimulation with cytokines significantly altered chromatin accessibility (log fold change > 1, FDR < 0.05) genome wide and co-stimulation with (TNF + IL-4) resulted in a distinct pattern of chromatin accessibility relative to stimulation with either cytokine alone (Extended Data Fig. 4b). We first examined regulation of chromatin accessibility by visualizing gene tracks at representative genes from each of the 5 gene clusters defined in Fig. 2c under the 4 stimulation conditions at 24 hr (Fig. 3a). Regulation of ATAC-seq peaks generally followed the pattern of gene expression, notably synergistic induction of open chromatin regions (OCRs) at the *CCL17* locus, and IL-4-mediated suppression of chromatin accessibility at the *CXCL10* locus. We then combined analysis of transcription factor expression (Fig. 3b) and computation of transcription factor activity score at OCRs using ChromVAR (Fig. 3c and Extended Data Fig. 4c) to test how differential regulation of TFs by cytokines is related to chromatin accessibility. Expression of the majority of IL-4-induced TFs, such as IRF4, BATF3 (which binds AP-1 sites) and EGR2, were superinduced by TNF, whereas TNF-induced factors such as IRF1 or STAT1 were suppressed by IL-4 (Fig. 3b). ChromVAR showed that STAT6 activity was increased by IL-4, whereas TNF increased NF-κB, AP-1 and CEBP activity. Under conditions of costimulation, STAT6 and NF-κB p65 activity were preserved, while AP-1 and CEBP activity was attenuated relative to that induced by TNF alone. Concomitant STAT6 and NF-κB p65 activity after co-stimulation was corroborated by analysis of signal enrichment around specific motifs using ChromVAR (Fig. 3d and Extended Data Fig. 4d). The overall pattern of additive STAT6 and NF-κB p65 binding, and attenuation of CEBP and AP-1 binding, upon co-stimulation with TNF + IL-4 was confirmed by an alternative computational strategy that detects ATAC-seq footprints using TOBIAS (Extended Data Fig. 4e). Together, the results suggest cooperation of STAT6 and NF-κB p65 under co-stimulation conditions contributes to synergistic gene induction, whereas IL-4 attenuates TNF-induced CEBP and AP-1 activity, and alters the pattern of expression of AP-1 proteins, to modulate the TNF response. These findings are in line with our observation of NF-κB and Stat6 motif enrichment in ATAC-seq analysis of mouse skin wound macrophages.

**Fig. 3.**
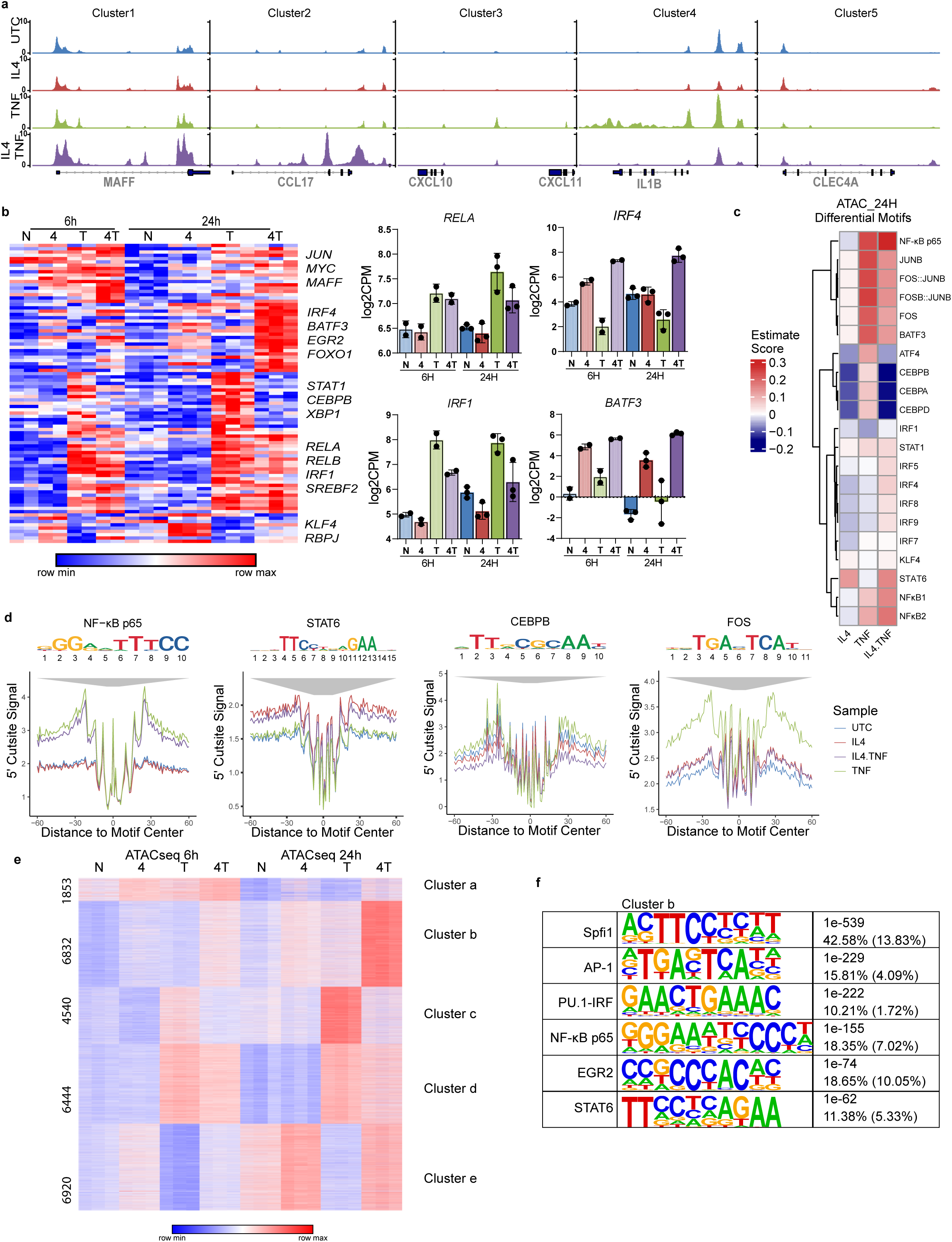
Regulation of chromatin accessibility and transcription factor activity. **a**, Representative Gviz tracks of selected genes in each of the clusters defined in Fig. 2. **b**, Left, Heatmap depicting expression of 91 transcription factors in the clusters defined in Fig. 2c. Right, Examples of TF gene expression. Log transformed counts per million (Log2CPM) are presented as the mean and SD of two (6 hour) or three (24 hour) individual donors per group quantified by RNA-seq. **c**, Heatmap of differentially enriched motifs at 24 hours comparing treatments relative to untreated controls from signal enrichment around specific motif analysis using ChromVAR. **d**, The signal enrichment profiles of NF-κB p65, STAT6, CEBPB and FOS at 24 hours using ChromVAR. **e**, K-means (K=5) clustering of differentially up-regulated peaks (log2 fold change > 1 and FDR < 0.05) in any pairwise comparison relative to untreated control. Biological replicates are shown in separate columns for each sample (left panel: 6h, n=3, right panel: 24h, n=3). **f**, The most significantly enriched transcription factor (TF) motifs identified by *de novo* motif analysis using HOMER in differentially regulated peaks in Cluster b from Fig. 3e.

The signal enrichment around specific motifs analysis described above was applied to the entire set of differentially regulated ATAC-seq peaks, and we reasoned that analysis of subsets of peaks regulated in a distinct manner by cytokines would be further informative. Therefore, we selected all up-regulated ATAC-seq peaks at 6 hours or 24 hours (log2 fold change > 1 and FDR < 0.05 compared to untreated controls) and clustered ATAC-seq peaks according to the pattern of regulation by cytokines (Fig. 3e). This analysis revealed 5 clusters of peaks, similar in pattern of regulation to the 5 gene clusters defined in Fig. 2, including peaks cooperatively induced by TNF and IL-4, and TNF-inducible peaks that were suppressed by IL-4. HOMER *de novo* motif analysis showed strong enrichment of both NF-κB-p65 and STAT6 motifs in peaks that were cooperatively induced by TNF and IL-4 (Fig. 3f). In contrast, TNF-induced peaks that were suppressed by IL-4 were enriched in AP-1, CEBP and IRF1 motifs (Extended Data Fig. 4f). These results reinforce the ChromVAR and TOBIAS analyses and support that cooperation between STAT6 and NF-κB p65 may contribute to synergistic gene induction, and suggest that IL-4 may target IRF1 in addition to AP-1 and CEBP to attenuate TNF-induced gene expression.

### IL-4 exploits TNF-induced NF-κB signaling to enhance its signature gene responses

The data suggesting cooperation of STAT6 and NF-κB p65 in driving synergistic gene induction led us to hypothesize that IL-4 might leverage TNF-mediated NF-κB responses to augment and expand its gene expression signature. We addressed this possibility by using ChIP-seq and CUT&RUN ^39^ to obtain genomic profiles of STAT6 and NF-κB p65 subunit binding, and of histone 3 lysine 27 acetylation (H3K27-Ac), which serves as a marker of regulatory element activation (Fig. 4a). In line with the results shown in Figure 3, the majority of IL-4-induced STAT6 peaks and of TNF-induced NF-κB p65 peaks were preserved under conditions of co-stimulation (Fig. 4a and Extended Data Fig. 5a, b). In Fig. 4a, left panels, the 10,750 total TNF-induced NF-κB p65 peaks have been segregated into peaks decreased by IL-4 (n = 2908, 27%), peaks unchanged by IL-4 (n = 6922, 64%) and peaks increased by IL-4 (n = 920, 9%). Interestingly, the NF-κB p65 peaks that were suppressed by IL-4 were associated with genes whose induction was suppressed by IL-4, such as *IL1A* and *IL1B*. We observed a substantial colocalization of STAT6 with NF-κB p65 binding at this set of genomic regions (Fig. 4a, compare middle and left panels, and Extended Data Fig. 5c). A subset of genomic regions exhibited increased binding of NF-κB p65 and STAT6 under co-stimulation conditions (Fig. 4a, bottom panels). Genomic regions with co-binding of NF-κB p65 and STAT6, especially those with increased binding of these transcription factors under co-stimulation conditions, were associated with synergistically induced genes such as *ALOX15*, *CCL22*, *BATF3* and *CCL17*. Examples of STAT6 and NF-κB p65 colocalization at synergistically induced genes are provided in gene tracks depicted in Fig. 4b. These include peaks that were only detected under co-stimulation conditions (Fig. 4b). In support of cooperative activity, peaks with co-binding of STAT6 and NF-κB p65 displayed increased levels of H3K27Ac signals (Fig. 4a, c). Additionally, STAT6 and NF-κB p65 binding was enriched in super-enhancers (Extended Data Fig. 5d), which are associated with (TNF + IL-4)-induced synergy genes as defined in Fig. 2e and listed in Supplementary Table 1 (Fig. 4d). To further examine the association of STAT6 and NF-κB p65 binding with gene expression, we separated genes into four categories based on their transcription factor binding patterns, as follows: genes with only NF-κB p65 peaks, genes with only STAT6 peaks, genes with NF-κB-p65 and STAT6 co-binding, and genes with no binding of these two transcription factors. The Empirical Cumulative Distribution Function (ECDF) Plot with Kolmogorov-Smirnov test for statistically significant differences in the relative expression of these genes showed that NF-κB p65 and STAT6 co-binding highly significantly up-regulated gene expression levels relative to genes with no binding or single transcription factor binding (Fig. 4e and Extended Data Fig. 5e). Finally, *de novo* motif analysis of STAT6-NF-κB p65 co-binding sites using HOMER showed enrichment of AP-1 motifs in addition to the expected NF-κB p65, STAT6 and PU.1 motifs, suggesting a possible role for a subset of AP-1 motif-binding transcription factors in the establishment and/or function of these regulatory elements (Extended Data Fig. 5f). Overall, the data support cooperation between STAT6 and NF-κB p65 as a mechanism by which TNF amplifies and extends the IL-4 response.

**Fig. 4.**
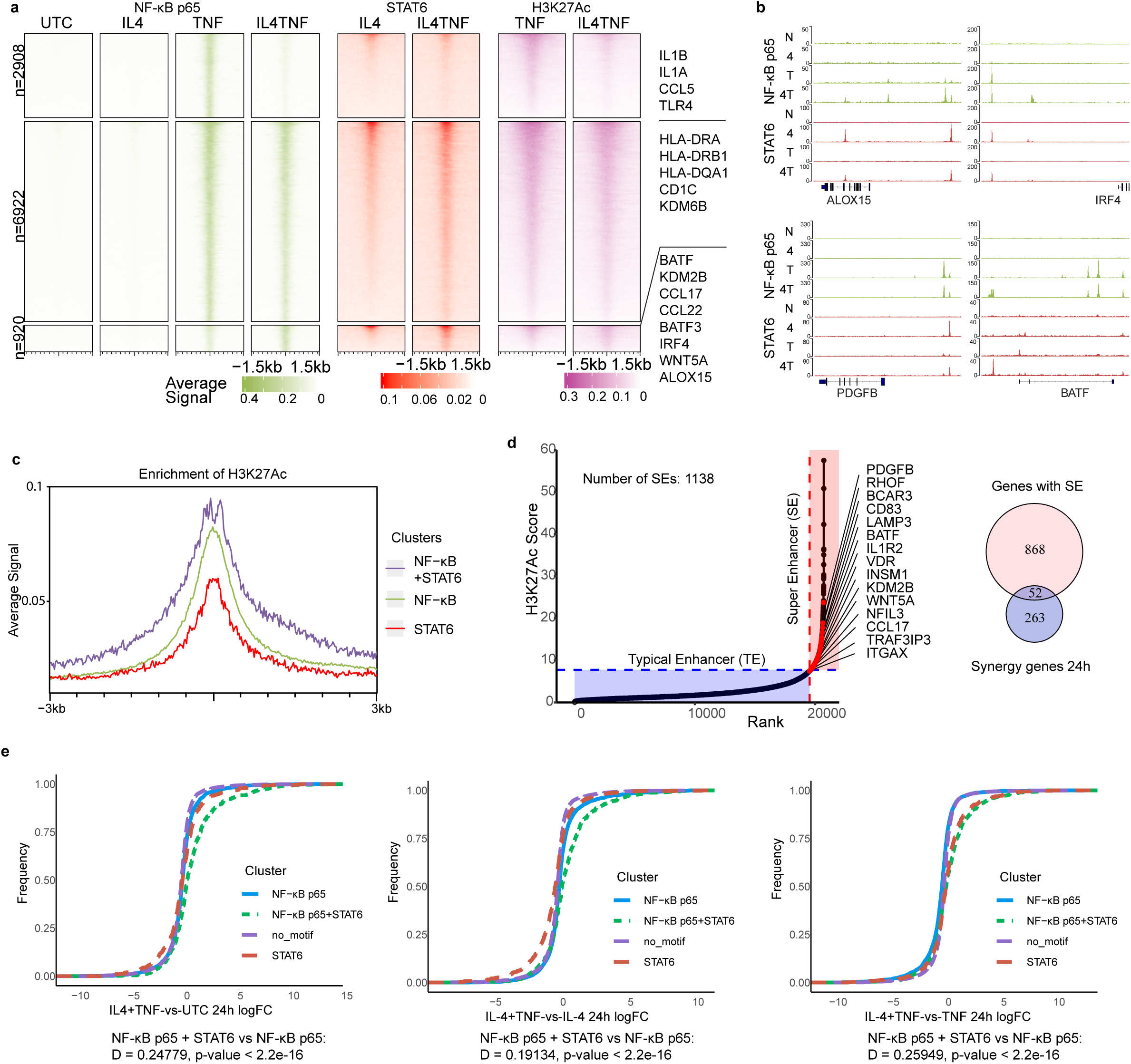
Cooperation between NF-κB and STAT6 to enhance gene responses. **a**, CUT&RUN and ChIP-seq analysis of NF-κB p65, STAT6 and H3K27Ac in human monocytes treated with indicated cytokines for 24 hours. Genomic regions surrounding NF-kB p65 binding peaks significantly induced by TNF or IL-4+TNF (FDR < 0.05) were grouped into three clusters using K-means (K=3) and the same genomic regions are depicted along the y-axis in all panels. Data are presented as normalized signal density ± 1.5kb around peak centers. Color scale represents the depth-normalized counts of peaks. NF-κB p65: n=4; STAT6: n=3; H3K27Ac: n=2. **b**, Representative Gviz tracks of synergy gene loci (*ALOX15, PDGFB*, *IRF4* and *BATF*). **c**, Average score of H3K27Ac enrichment at CUT&RUN peaks showing binding by NF-κB p65 alone, STAT6 alone or both NF-κB p65 and STAT6. **d**, Left, signal intensities of H3K27Ac peaks in IL-4 and TNF co-stimulation condition were counted and ranked, super enhancers are defined as regions whose intensity is above the tangent of slope 1. Selected synergy genes associated with super enhancers are highlighted. Right, Venn diagram of the overlap between synergy genes and genes with super enhancers under co-stimulation conditions. **e**, ECDF plot with Kolmogorov-Smirnov statistical test. The x-axis displays the log2 fold change (log2FC) of gene expression, representing the magnitude and direction of gene expression changes under different conditions. The y-axis quantifies the cumulative frequency of genes at each log2FC level. Four categories of genes are plotted: genes with only NF-κB p65 peaks, with only STAT6 peaks, with NF-κB-p65 and STAT6 co-binding, and genes with no binding of these two transcription factors. The D value of Kolmogorov-Smirnov Test indicates the differences of the log2FC between selected comparison (genes with NF-κB-p65 and STAT6 co-binding versus genes with only NF-κB p65 peaks). Values of p < 0.05 were considered statistically significant.

### NF-κB Signaling Augments IL-4-induced Gene Expression

We next inhibited NF-κB signaling to directly test the role of NF-κB in induction of synergy genes that co-bound STAT6 and NF-κB p65, such as *ALOX15*, *CCL17*, and *IRF4* (as depicted in Fig. 4b). Three different approaches using an IKKα/β inhibitor, an IκBα phosphorylation inhibitor and adenoviral-mediated overexpression of an IκBα super repressor yielded similar results: synergy genes *ALOX15*, *CCL17* and *IRF4* were strongly dependent on NF-κB signaling under costimulation conditions (Fig. 5a - c). In contrast, genes that were induced by IL-4 alone, such as *SOCS1* and *MRC1*, were minimally affected by NF-κB inhibition (Fig. 5d). Immunoblot results indicated that suppression of synergy gene expression was not due to reduced STAT6 activity (Fig. 5e). We then used ChIP-qPCR to test whether inhibition of NF-κB signaling affected STAT6 recruitment to target genes. STAT6 recruitment to ALOX15 and SOCS1 promoters was modestly decreased by NF-κB pathway inhibition after IL-4 stimulation or co-stimulation with TNF and IL-4 (Fig. 5f) which is insufficient to explain the strong inhibition of gene induction and suggests that NF-κB primarily contributes to gene induction directly rather than by regulating STAT6.

**Fig. 5.**
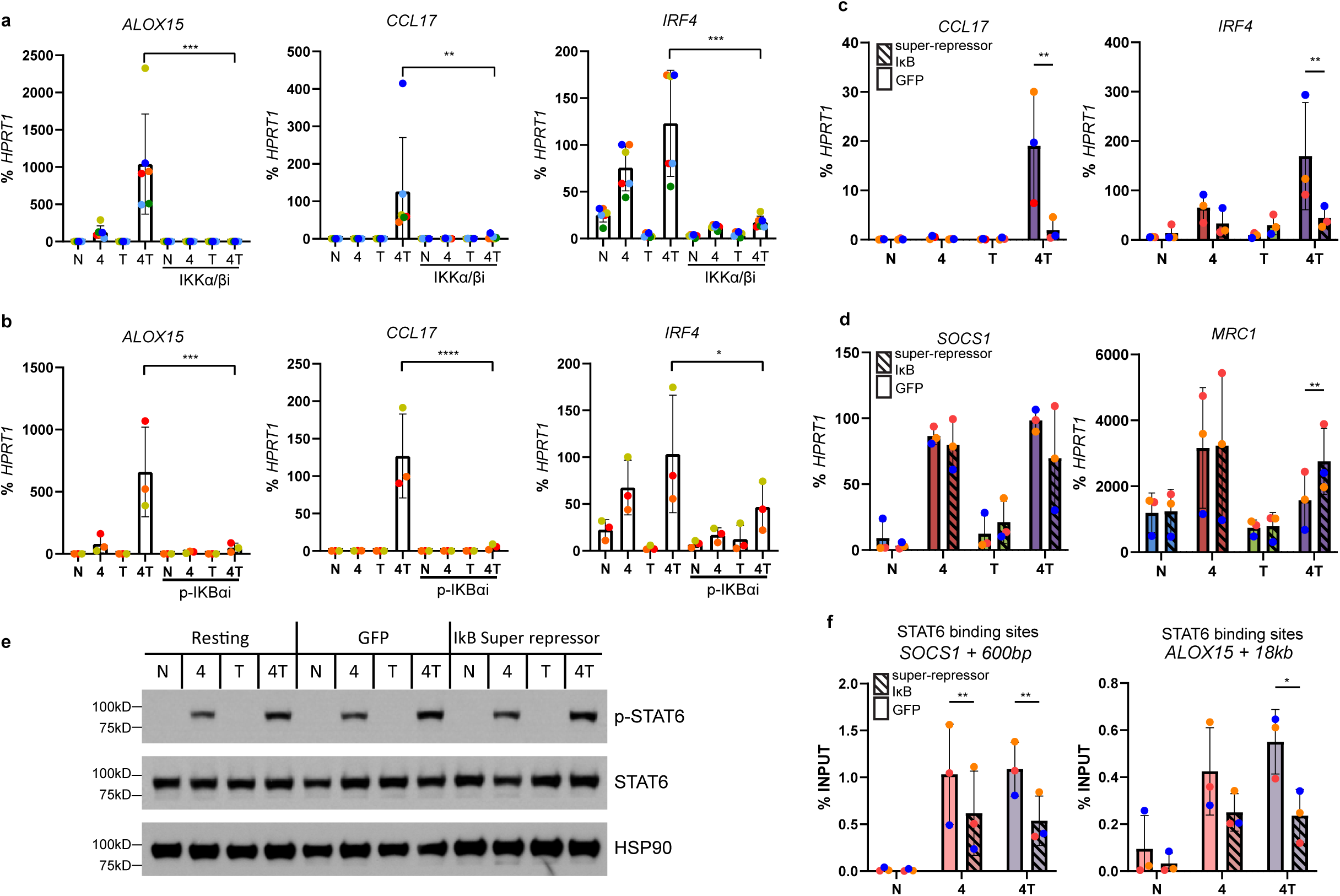
NF-κB Signaling Augments IL-4-induced Gene Expression. **a**, **b**, mRNA of indicated genes measured by RT-qPCR and normalized relative to *HPRT1* mRNA in cells pre-treated with IKKα/β inhibitor (10 µM) (**a**) or IκBα phosphorylation inhibitor (20 µM) (**b**) for 30 min and then stimulated with indicated cytokines for 6h. n = 6 (B) or 3 (C). **c**, **d,** Human CD14^+^ monocytes were cultured for 5 days with M-CSF and then transduced with adenoviral particles encoding eGFP or IκBα super-repressor. Cells were rested overnight after transduction and stimulated with indicated cytokines for 3 hours. mRNA of indicated genes measured by RT-qPCR and normalized relative to *HPRT1.* Data are presented as mean +/− SD, n = 3. **e**, Western blot of tyrosine-phosphorylated STAT6 in human macrophages transduced with adenoviral particles as described in **c**. One representative blot from three independent experiments is shown. HSP90 serves as a loading control. **f,** ChIP-qPCR analysis of genomic regions of human macrophages transduced with adenoviral particles encoding eGFP or IκB super-repressor and stimulated with indicated cytokines for 3 hours.

### Cooperative erasure of the negative H3K27me3 histone mark at synergy genes

TNF and IL-4 synergistically activated *KDM6B* (Fig. 6a), which encodes a Jumonji C domain-containing lysine demethylase that erases the negative histone mark H3K27me3 to facilitate transcription. The *KDM6B* locus co-bound STAT6 and NF-κB p65 (Fig. 6b) and induction of *KDM6B* mRNA after co-stimulation was dependent on NF-κB signaling (Fig. 6c), which raises the possibility that the TNF-NF-κB axis may also augment IL-4-induced gene expression via KDM6B-mediated loss of H3K27me3. Indeed, TNF co-stimulation potentiated IL-4-mediated decreases in H3K27me3 at the *ALOX15* and *IRF4* promoters (Fig. 6d). We then used CUT&RUN to obtain the genomic profile of H3K27me3, and also of the H3K4me3 which marks active promoters and is associated with transcription. In line with the ChIP-qPCR data, genes that were strongly induced after TNF + IL-4 co-stimulation (corresponding to clusters 1 and 2 in Fig. 2c) showed baseline H3K27me3 surrounding the transcription start site (Fig. 6e), which was particularly prominent for a subset of genes in cluster 2, where the strongest co-stimulation of gene expression was observed at the 24 hr timepoint. These genes showed a strong decrease in H3K27me3 in the (TNF + IL-4) condition (Fig. 6e, f); in contrast H3K4me3 increased in parallel with gene expression as expected. Representative gene tracks showing loss of H3K27me3 and concomitant gain of H3K4me3 at the *ALOX15* and *IRF4* loci are shown in Fig. 6g; in contrast to these synergy genes, genes induced by TNF alone (*CCL5* and *IL1B*) or IL-4 alone (*CISH*, *SOCS1*) showed minimal H3K27me3 levels at baseline and these did not appreciably change after cytokine stimulation. A functional role for the KDM6B-H3K27me3 axis in restraining expression of synergy genes was supported by decreased induction of these genes when KDM6B was inhibited to maintain basal H3K27me3 levels (Fig. 6h). Overall, these results show that TNF signaling via NF-κB also promotes induction of synergy genes after TNF + IL-4 co-stimulation by an indirect mechanism, via induction of KDM6B and erasure of the negative histone mark H3K27me3.

**Fig. 6.**
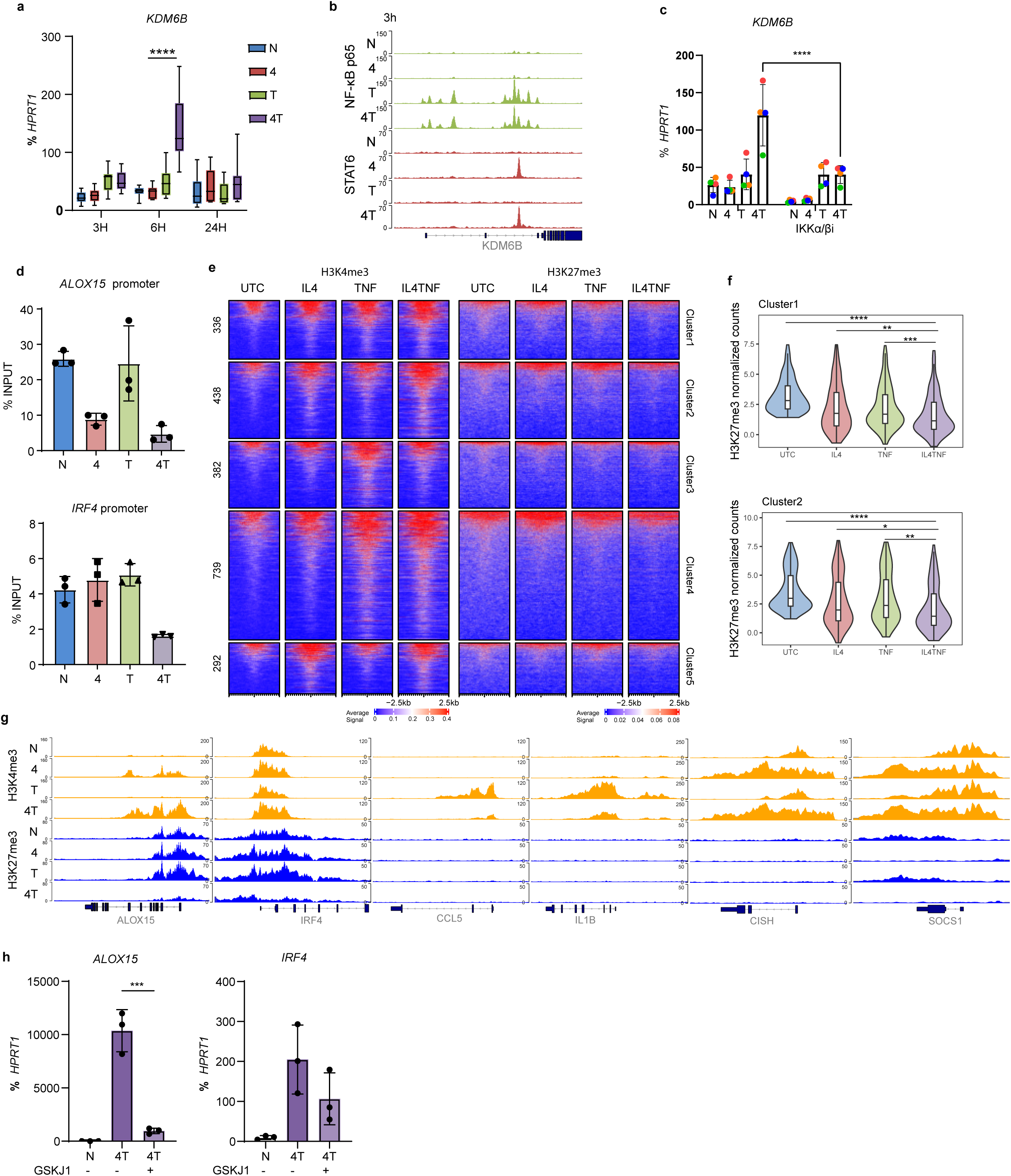
Cooperative erasure of the negative H3K27me3 histone mark at synergy genes. **a**, mRNA of *KDM6B* measured by RT-qPCR and normalized relative to HPRT1 mRNA in human monocytes treated with indicated cytokines for 3, 6 and 24 hours. Same samples as in Fig. 2b, n = 9. One-way ANOVA, ****p value < 0.0001. **b**, Representative Gviz tracks showing *KDM6B* locus with NF-κB-p65 and STAT6 binding at 3 hours after indicated stimulation. **c**, mRNA of *KDM6B* measured by qPCR and normalized relative to *HPRT1* mRNA in cells pre-treated with IKKα/β inhibitor (10 µM) for 30 min and stimulated with indicated cytokines for 6 hours. One-way ANOVA, ****p value < 0.0001. n=4. **d**, ChIP-qPCR analysis of H3K27me3 at *ALOX15* and *IRF4* loci of human monocytes at 24 hours after indicated stimulation. n = 3. **e**, CUT&RUN analysis of H3K27me3 and H3K4me3 on differentially down-regulated (H3K27me3) up-regulated or (H3K4me3) peaks associated with genes in clusters identified in Fig. 2c. Data are presented as normalized signal density ± 2.5kb around peak centers. Color scale represents the depth-normalized counts of peaks. n = 3. **f**, The violin plots represent cumulative values of normalized H3K27me3 signals at genes in cluster 1 and cluster 2 as shown in panel **e** under indicated conditions. Pairwise Wilcox test, *p < 0.05, **p < 0.01, ***p < 0.001, ****p < 0.0001. **g**, Representative Gviz tracks showing H3K3me3 and H3K27me3 signals at the indicated gene loci. **h**, mRNA of *ALOX15* and *IRF4* measured by RT-qPCR and normalized relative to *HPRT1* mRNA in cells pre-treated with GSKJ1 (50 µM) and stimulated with indicated cytokines for 6 hours. One-way ANOVA, **p < 0.01. Error bars represent SD.

### IL-4 targets IRF1-binding genomic elements to suppresses TNF-induced interferon responses

We wished to further investigate how IL-4 suppressed components of the TNF response. We confirmed that IL-4 effectively suppressed the TNF-induced IFN response using gene set enrichment analysis (GSEA) (Fig. 7a). As the TNF-induced IFN response is IRF1-dependent ^27^, and ATAC-seq peaks induced by TNF whose accessibility was suppressed by IL-4 were enriched in the IRF1 binding motif (Extended Data Fig. 4f), we tested whether IL-4 suppresses IRF1. We used CUT&RUN to obtain a genomic profile of IRF1 binding (Extended Data Fig. 6a) and found that IRF1 peaks were associated with IFN response genes (Extended Data Fig. 6b), and that IL-4 broadly suppressed IRF1 binding across the genome (Fig. 7b). Representative gene tracks showing IRF1 binding at the ISGs *CXCL10* and *ISG15* are shown in Fig. 7c. Decreased genomic binding by IRF1 was explained at least in part by IL-4-mediated suppression of IRF1 mRNA and nuclear IRF1 protein (Fig. 3b and Extended Data Fig. 6c). These results, together with the known dependence of the TNF-induced IFN response on IRF1, show that IL-4 suppresses the TNF-induced IFN response at least in part by inhibiting IRF1. However, add-back of IRF1 via adenoviral-mediated transduction did not reverse suppression of ISGs by IL-4 (Fig. 7d), suggesting that IL-4 suppresses additional components that induce the TNF-mediated IFN response. *De novo* motif analysis using HOMER showed enrichment of RUNX1, CEBP, NF-κB p65 and AP-1 motifs under IRF1 peaks (Fig. 7e), suggesting that IL-4 may also target binding of these TFs at key IRF1-binding elements important for the IFN response. As IRFs and NF-κB p65 can bind cooperatively at some genomic elements, we first tested the effects of IL-4 on NF-κB p65 binding at IRF1 peaks. Although NF-κB p65 binding was observed at the majority of IRF1 peaks, co-treatment with IL-4 that strongly suppressed IRF1 minimally affected NF-κB p65 binding (Fig. 7f). Furthermore, when we subsetted IRF1 binding sites into two categories: with or without NF-κB p65 co-binding, IL-4 suppressed IRF1 binding comparably at both types of genomic elements (Fig. 7g, NF-κB p65 binding is plotted as a control). Investigation of other transcription factors that can bind to genomic elements corresponding to IRF1 leaks, as shown in Fig. 7e, revealed that IL-4 strongly suppressed TNF-induced expression of CEBPβ, a pioneer transcription factor that opens chromatin (Fig. 7h). These results suggest that IL-4 selectively inhibits the TNF-induced IFN response by targeting a set of genomic elements characterized by IRF1 and CEBP binding.

**Fig. 7.**
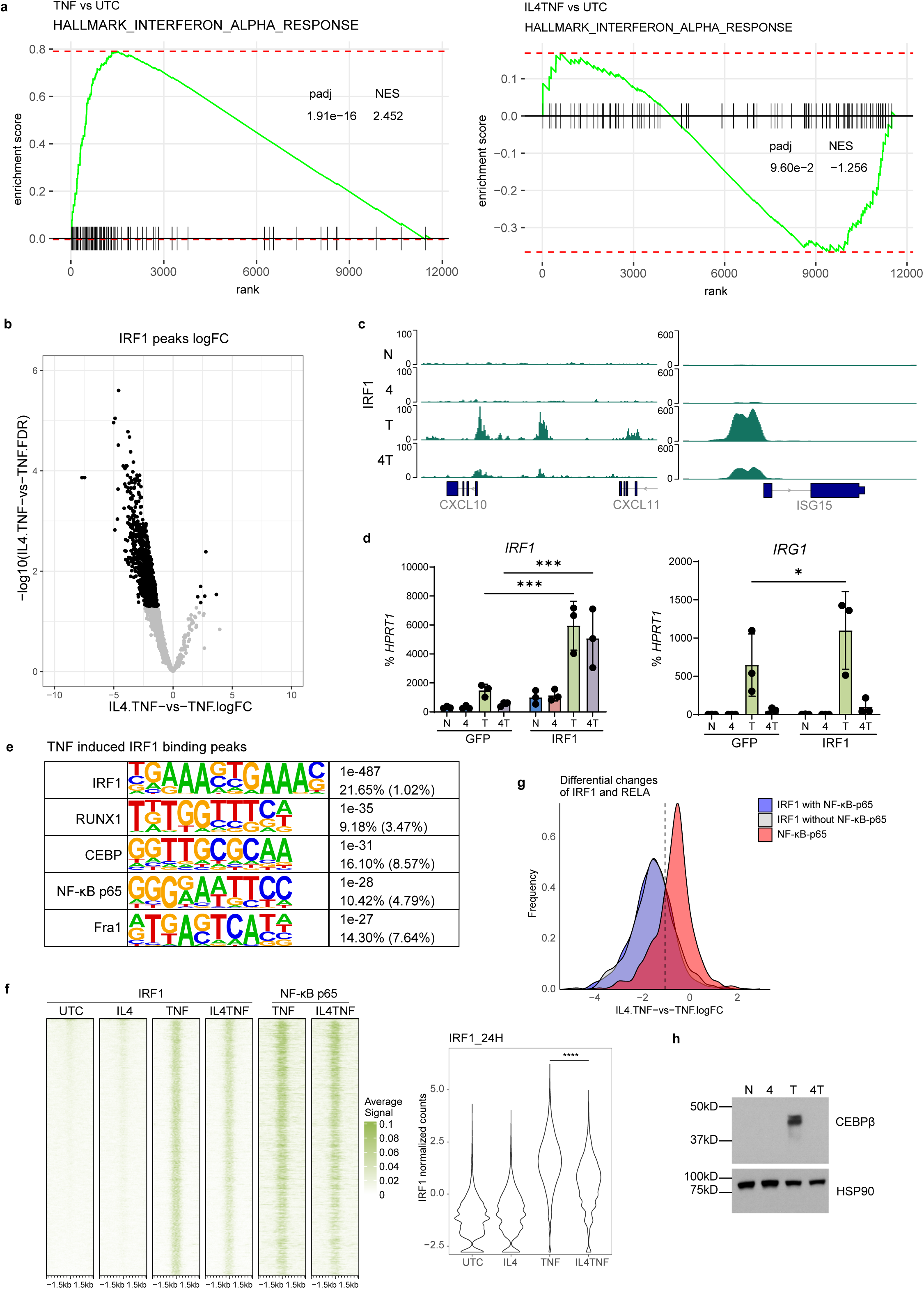
IL-4 targets IRF1-binding genomic elements to suppresses TNF-induced interferon responses. **a**, GSEA results of ranked differentially expressed genes from RNAseq data in Fig. 2 (n = 3) using the HALLMARK_INTERFERON_ALPHA_RESPONSE gene set. The Normalized Enrichment Score (NES) provide a normalized measure of the degree to which a set of genes is overrepresented at the top or bottom of a ranked list of genes. Values of adjusted p value < 0.05 were considered statistically significant. **b**, Volcano plots of statistical significance (-log10 FDR, y axis) plotted against log2 ratio of differential IRF1 binding based on IRF1 CUT&RUN data showing differentially induced (right) or suppressed (left) peaks after combined treatment with IL4 and TNF compared to TNF treated monocytes. Significantly differentially expressed peaks (log2 fold change > | 1|; FDR < 0.05) (black), non-significant peaks (grey). **c**, Representative Gviz tracks showing IRF1 binding at the indicated loci *CXCL10* and *ISG15*. **d**, Human CD14^+^ monocytes were first cultured for 5 days with M-CSF, then transduced with adenoviral particles encoding eGFP or IRF1. Cells were rested for 24 hours after transduction and stimulated with indicated cytokines for 6 hours. mRNA of indicated genes measured by RT-qPCR and normalized relative to *HPRT1.* Data are depicted as mean ± SD. **e**, *De novo* motif analysis using HOMER of TNF-induced IRF1 binding sites after 24 hours stimulation. **f,** Left Panel: CUT&RUN of IRF1 and NF-κB p65 binding in human monocytes treated with indicated cytokines for 24 hours. The peaks depicted (y-axis) correspond to all significantly induced IRF1 peaks (fold change > 1, FDR < 0.05). Data are presented as normalized signal density ± 1.5kb around peak centers. Color scale represents the depth-normalized counts of peaks. Right panel: Violin plots indicate normalized log2CPM counts of IRF1 binding. Mann-Whitney U Test, ****p < 0.0001. **g**, Histogram plot of IRF1 peaks with/without NF-κB p65 co-localization and NF-κB p65 peaks as control, signals are displayed as log2 fold-change between IL4+TNF and TNF conditions. The y-axis shows the cumulative frequency of peaks at each log2FC level. **h**, Immunoblot of CEBPβ and HSP90 using whole cell lysates 24 hours after indicated stimulations. One representative blot from six independent experiments is shown. HSP90 serves as loading control. (**b**-**f**) IRF1 CUT&RUN, n = 3.

## DISCUSSION

In myeloid cells IL-4 elicits a core STAT6-mediated transcriptional response whose modulation and expansion by tissue factors is important for host defense and tissue homeostasis and repair ^20, 21^. Mechanisms by which distinct factors expand IL-4 responses are not known, and the effects of inflammatory factors on IL-4 signaling are not well characterized but are assumed to be mostly antagonistic based on prior work with IFN-γ and the current paradigm of mutual suppression by cytokines with opposing biological functions ^15, 40^. Surprisingly, in this study we found that instead of antagonizing IL-4 activity, the potent inflammatory cytokine TNF greatly augmented and expanded the IL-4 transcriptome, including genes important in immune cell activation, metabolism, and chemoattraction. Reciprocally, IL-4 selectively suppressed the TNF-induced IFN response while leaving inflammatory gene induction mostly intact. Crosstalk between TNF and IL-4 was mediated by epigenetic chromatin-mediated mechanisms that were associated with cooperation between NF-κB p65 and STAT6, erasure of H3K27me3, and selective inhibition of IRF1. Induction of TNF-IL-4 synergy genes was observed in a subset of macrophages *in vivo* in a skin wound model where TNF and IL-4 are co-expressed. These results identify mechanisms that expand the IL-4 response and mediate the crosstalk between opposing cytokines that is required for an effective and focused inflammatory response that avoids IFN-associated toxicity.

The transcriptional profile and biology of TNF + IL-4 co-stimulated monocytes were quite distinct from that of monocytes stimulated with either cytokine alone. TNF not only amplified the IL-4 response but extended it through induction of synergy genes and *de novo* activation of genes that were not induced by either cytokine alone. Expression of these synergy genes alters the monocyte state by engaging pathways important in immune activation and in cholesterol, fatty acid and steroid biosynthesis. Changes in lipid metabolism are associated with an M2-like and tissue repair phenotype, and with modulation of IFN responses ^41, 42^. Costimulation also induced expression of genes that affect interactions of monocytes with other cell types: presentation of protein and lipid antigens to T cells, the PD-1 pathway that inhibits T cells, and chemoattractants. Activation of the PD-1 pathway and induction of chemokines such as CCL17 and CCL22 that recruit Tregs and type 2 lymphocytes supports that TNF costimulation promotes homeostatic and tissue-trophic functions of IL-4. Accordingly, co-stimulation induced expression of growth factors such as *TGFA*, *WNT5A*, *INHBA* and *PDGFB*, which promote the proliferative phase of wound healing. Interestingly and in line with a trophic phenotype, macrophages that express aspects of this costimulatory program were present in skin wounds at a time when both TNF and IL-4 are expressed and wounds are transitioning from an inflammatory to a proliferative and reparative phase. The role of TNF in skin wound healing is complex and context- and time-dependent. For example, loss of TNF signaling can result in increased angiogenesis ^43^, augmented fibroblast activation and enhanced scar tissue formation ^44^, while injection of TNF has been shown to accelerate the wound healing process ^45^. Selective suppression of the TNF-induced IFN response may contribute to the physiological reduction of inflammation required for the transition to the reparative phase. Given the preservation of the TNF-driven NF-κB-mediated response and the induction of antigen-presenting molecules and immune mediators, it is possible that in some contexts IL-4-TNF crosstalk can contribute to excessive inflammation and pathology, for example in conditions like asthma where both cytokines are highly expressed in a sustained manner.

The mechanisms of synergistic gene induction involved cooperation between core transcription factors in TNF and IL-4 signaling, NF-κB p65 and STAT6, respectively. One direct mechanism involved concomitant binding of NF-κB p65 and STAT6 to genomic elements including superenhancers, which was associated with increased chromatin accessibility, histone acetylation, and transcription. This contrasts with crosstalk between IL-4 and IFN-γ, which is mostly antagonistic and mediated by suppression of auxiliary transcription factors, while core STAT6 and STAT1 signaling remains mostly unchanged ^15^. A second and indirect mechanism involved synergistic induction by IL-4 and TNF of KDM6B, which broadly erased the negative histone mark H3K27me3 surrounding the TSS at synergistic target genes. This extends previous work implicating KDM6B in induction of IRF4 by IL-4 and GM-CSF ^14, 18, 19, 46^ by showing erasure of H3K27me3 at multiple additional genes, including key transcription factors like BATF3 and EGR2. Interestingly, induction of synergy genes was associated with a low baseline and strong induction of positive promoter mark H3K4me3, which is reminiscent of “bivalent” genes in embryonic stem cells ^47, 48, 49^. Notably, this coordinate induction of H3K4me3 and erasure of H3K27me3 is found not only in key transcription factor genes (such as *IRF4*, *BATF3*, and *EGR2*) but also in *PDGFB* and *TGFA*, which stimulate fibroblast and keratinocyte proliferation and the production of collagen and extracellular matrix proteins and play an important role in tissue repair. In line with the findings in human monocyte promoters, we found opening of chromatin at synergy gene promoters in mouse skin wound F4/80-low macrophages ^50, 51^. Promoters of most inducible genes in macrophages typically exhibit open chromatin at baseline. In contrast, we have newly identified a gene set with low baseline expression and inaccessible promoter chromatin that is massively induced by TNF and IL-4 in association with *de novo* opening of gene promoters.

In contrast to augmentation of the IL-4 response by TNF, IL-4 selectively suppressed the TNF-induced IFN response by targeting IRF1. IRF1 is not only required for induction of *IFNB* and the ensuing autocrine loop, but also bound to multiple ISGs, half of the time in association with NF-κB. IL-4 strongly suppressed IRF1 binding with a minimal effect on NF-κB p65 cobinding including at ISGs, indicating that NF-κB p65 binding at ISGs is not dependent on IRF1 and is insufficient to activate transcription. Indeed, IL-4 had minimal effects on TNF-induced NF-κB p65 binding genome-wide, which is in accord with limited suppression of pro-inflammatory genes such as *IL1B* that rely more on the canonical NF-κB signaling pathway. This stands in striking contrast to IL-4-mediated regulation of the LPS response, where IL-4 expands the NF-κB p65 cistrome and does not affect the IFN response ^11^. These differences are most likely explained by different signaling pathways utilized by TLR4 and TNF receptors to activate NF-κB and IFN responses. Activation of IFN responses by TLR4 is not dependent on IRF1 but instead is mediated by TRIF-IRF3 signaling ^38^. Thus, the preferential utilization of IRF1 for induction of IFN responses by TNF confers sensitivity to inhibition by IL-4.

In summary, our research has revealed the crucial role of the transcription factor NF-κB p65 in mediating IL-4’s effects when cells are co-treated with TNF, shedding light on their intricate interplay at the chromatin level. Importantly, this utilization of NF-κB is highly selective, as we observed a substantial suppression of IRF1, another crucial transcription factor induced during late-phase TNF treatment, while NF-κB p65 signals remained predominantly unaffected. This regulatory mechanism extends its influence beyond the scope of IL-4 signature genes, impacting genes associated with diverse functions such as antigen presentation, steroid metabolism, and chemoattractant activity. These findings imply that monocytes and macrophages displaying this distinctive phenotype may be pivotal in coordinating the transition between different stages of wound healing. This research has implications for modulating the interplay of cytokines with opposing functions to promote tissue repair and a return to homeostasis, while avoiding pathology associated with excessive inflammation.

## METHODS

### Study design

The study focused on the epigenomic regulation of inflammatory responses in monocytes under the co-stimulation of TNF and IL-4, and their impact on inflammation and tissue repair. CD14^+^ monocytes were purified from healthy donors at the New York Blood Center (Long Island City, NY). To investigate transcriptional and epigenetic changes, we conducted RNA-seq, ATAC-seq, ChIP-seq, and CUT&RUN analyses on co-stimulated monocytes. We also examined the phenotypes of mouse monocytes/macrophages in skin wounds four days post-injury, the time when TNF and IL-4 are co-expressed, utilizing multiparameter flow cytometry sorting and RT-qPCR analysis. Additionally, a re-analysis of the single-cell study (GSE142471) was undertaken to evaluate monocyte and macrophage subpopulations during the wound healing process. The number of biological replicates is detailed in the figure legends. Neither blinding nor randomization was employed in this study. For the genome-wide sequencing, experiments donors were screened for lack of baseline activation, and for engagement of the TNF-IFN autocrine loop, as assessed by TNF-induced CXCL10 expression.

### Mice

Animal experiments were approved by the Weill Cornell Medicine IACUC Committee. C57BL/6J mice (Strain #:000664) and *TNF^−/−^* mice (Strain #: 003008) at 6 to 8 weeks old were purchased from the Jackson Laboratories. All the mice were housed under specific pathogen-free conditions. Full-thickness dorsal skin wounds were created by excision 4-mm biopsy punches (Miltex) on either side of midline. Two wounds were generated in the upper back of mice for flow cytometry analysis of cellular proportions; 4 wounds were generated in the upper and lower back of mice for cell sorting and wound closure experiments. Prior to the surgery, age-matched, randomly assigned male mice were anesthetized with isoflurane and subcutaneous injections of buprenorphine (0.5 mg/kg) were given as analgesics. At the indicated time points, mice were euthanized, and wound tissues were collected with 8-mm biopsy punches (Miltex). For inhibition experiments, the vehicle ointment was prepared by mixing vaseline (Sigma) and sunflower oil (Sigma) in a 1:1 ratio (w/w) under sterile conditions. To produce the STAT6 inhibitor ointment, AS1517499 (STAT6 inhibitor) powder was incorporated into the vehicle ointment. The mixture was homogenized using a T25 digital disperser (IKA) at 12,000 rpm until uniform consistency was achieved. Either the vehicle or inhibitor ointment was applied directly to the wound using a disposable Calcium Alginate Tipped Applicator (Puritan Medical Products Co LLC) at final concentrations of 0.4 nmol/kg.

### Human cells

Deidentified human buffy coats were purchased from the New York Blood Center following a protocol approved by the Hospital for Special Surgery Institutional Review Board. Peripheral blood mononuclear cells (PBMCs) were isolated using density gradient centrifugation with Lymphoprep (Accurate Chemical) and monocytes were purified with anti-CD14 magnetic beads from PBMCs immediately after isolation as recommended by the manufacturer (Miltenyi Biotec). Monocytes were cultured overnight at 37°C, 5% CO_2_ in RPMI-1640 medium (Invitrogen) supplemented with 10% heat-inactivated defined FBS (HyClone Fisher), penicillin-streptomycin (Invitrogen), L-glutamine (Invitrogen) and 20 ng/ml human M-CSF (Peprotech). Then, the cells were treated as described in the figure legends.

### Analysis of RNA and protein

Total RNA was extracted with an RNeasy Mini Kit (QIAGEN) and was reverse-transcribed with a RevertAid RT Reverse Transcription Kit (Thermo Fisher Scientific). Real-time PCR was performed with Fast SYBR Green Master Mix and a 7500 Fast Real-time PCR system (Applied Biosystems). CT values of target genes were normalized to *HPRT1* for human and *Gapdh* for mouse cells, and are shown as percentage of *HPRT1* or *Gapdh* (100/2^^ΔCt^). The primer sequences for the quantitative RT-qPCR reactions are listed on Supplementary Table 2.

For protein analysis, cells were washed with cold PBS after specified treatments and harvested in SDS lysis buffer (2% SDS, 50mM Tris-HCl, pH 6.8, 10% (v/v) glycerol) supplemented with 1x phosSTOP EASYPACK and 1× EDTA-free complete protease inhibitor cocktail (Roche, Basel, Switzerland) and incubated for 15 min on ice. Then, cell debris was pelleted at 16,000 x *g* at 4°C for 10 min. The soluble protein fraction was mixed with 4× Laemmli Sample buffer (Bio-Rad) and 2-mercroptoehanol (BME) (Sigma-Aldrich). Samples for Western blotting were subjected to electrophoresis on 4–12% Bis-Tris gels (Invitrogen). Proteins were transferred to polyvinylidene difluoride membrane as previously reported.^52^ Membranes were blocked in 5% (w/v) Bovine Serum Albumin in TBS (20 mm Tris, 50 mm NaCl, pH 8.0) with 0.1% (v/v) Tween 20 (TBST) at room temperature for at least 1 h with shaking at 60 rpm. Membranes were then incubated with primary antibodies at 4 °C overnight with shaking at 60 rpm. Membranes were washed 3 times in TBST, then probed with anti-mouse or anti-rabbit IgG secondary antibodies conjugated to horseradish peroxidase (GE Healthcare, cat: NA9310V and NA9340V) diluted in TBST at room temperature for one hour with shaking at 60 rpm. Next, membranes were washed 3 times in TBST at room temperature for with shaking at 60 rpm. Antibody binding was detected using enhanced chemiluminescent substrates for horseradish peroxidase (HRP) (ECL Western blotting reagents (PerkinElmer, cat: NEL105001EA) or SuperSignal West Femto Maximum Sensitivity Substrate (Thermo Fisher Scientific, cat: 34095), according to the manufacturer’s instructions, and visualized using premium autoradiography film (Thomas Scientific, cat: E3018). To detect multiple proteins on the same experimental filter, membranes were cut horizontally based on the molecular size of the target proteins. For membranes that required probing twice or more using different primary antibodies, RestoreTM PLUS Western blotting stripping buffer (Thermo Fisher Scientific) was applied on the blots with shaking at 60 rpm for 15 min following previous development. The antibodies used were from Cell Signaling Technology: STAT6 (5397S), phosphor-STAT6 (9361S), C/EBPβ (3087S), HSP90α (8165S), IRF1 (8478S), from Sigma-Aldrich: α-Tubulin (T9026) and from Abcam: Lamin B1 (ab16048).

### RNA sequencing

Libraries for sequencing were prepared using extracted RNA and the NEBNext Ultra II RNA Library Prep Kit from New England Biolabs (NEB) following the manufacturer’s instructions. Quality of all RNA and library preparations was evaluated with BioAnalyser 2100 (Agilent). Libraries were sequenced by the Genomic Resources Core Facility at Weill Cornell Medicine using a HiSeq2500, 50-bp single-end reads at a depth of ∼20 - 40 million reads per sample. Read quality was assessed and adapters trimmed using FastQC and cutadapt. Reads were then mapped to the human genome (hg38) or mouse genome (mm10) and reads in exons were counted against Gencode v27 with STAR Aligner ^53^. Differential gene expression analysis was performed in R using edgeR ^54^. Only genes with expression levels exceeding 3 counts per million in at least one group were utilized for downstream analysis. Benjamini-Hochberg false discovery rate (FDR) procedure was used to correct for multiple testing. Heatmap with K-mean clustering was done using Morpheus web application. Heatmaps for each cluster were drawn using the R package pheatmap ^55^.

### Gene Set Overrepresentation and Enrichment Analysis

The overrepresentation and enrichment of terms from KEGG (https://www.genome.jp/kegg/), Reactome ^56^ (https://reactome.org), BioPlanet (https://tripod.nih.gov/bioplanet/), NCI (https://datascience.cancer.gov/resources/nci-data-catalog) and Gene Ontology (GO, http://geneontology.org) in a specific list of genes or ranked genes were investigated. The gene sets were obtained using R msigdbr package ^57^. The hypergeometric test was used to measure the degree of enrichment. The ORA was implemented in R clusterProfiler ^58^.

### ATAC sequencing

ATAC-seq libraries were prepared as described previously ^59^. Briefly 5×10^4^ cells were spun at 500 x *g* for 5 min at 4 °C, which was followed by a wash using 50 ml of cold PBS and centrifugation at 500 x *g* for 5 min. Cells were lysed using cold lysis buffer (10 mM Tris-HCl, pH 7.4, 10 mM NaCl, 3 mM MgCl2 and 0.1% IGEPAL CA-630). Immediately after lysis, nuclei were spun at 500 x *g* for 10 min in a refrigerated centrifuge. Immediately following the nuclei prep, the pellet was resuspended in the transposase reaction mix (25 µl 2× TD buffer, 2.5 µl transposase (Illumina) and 22.5 µl nuclease-free water). The transposition reaction was carried out for 30 min at 37 °C. Directly following transposition, the sample was purified using a MinElute PCR Purification kit. Then, we amplified library fragments using 1× NEB next PCR master mix and 1.25 M of custom Nextera PCR primers as previously described ^28^, using the following PCR conditions: 72 °C for 5 min; 98 °C for 30 s; and thermocycling at 98 °C for 10 s, 63 °C for 30 s and 72 °C for 1 min. The libraries were purified using a Qiagen PCR cleanup kit yielding a final library concentration of ∼30 nM in 20 µl. Libraries were amplified for a total of 10–13 cycles and were subjected to high-throughput sequencing at the Genomic Resources Core Facility at Weill Cornell Medicine using the Illumina HiSeq 4000 Sequencer (paired end reads) to at least 20 million reads/sample. ATAC-seq data was aligned to the genome using the same pipeline as ChIP-seq and CUT&RUN data (see below). Data for ATAC-seq experiments are from three independent experiments with different blood donors.

### ChIP and ChIP sequencing

Cells were crosslinked for 5 min at room temperature by the addition of fresh, methanol-free 16% formaldehyde (w/v) (Pierce) to the growth medium to a final concentration of 1%, followed by 5 min of quenching with 100 mM glycine. Cells were pelleted at 4 °C and washed with ice-cold PBS. The crosslinked cells were lysed with lysis buffer (50 mM HEPES-KOH, pH 7.5, 140 mM NaCl, 1 mM EDTA, 10% glycerol, 0.5% NP-40 and 0.25% Triton X-100) with protease inhibitors on ice for 10 min and were washed with washing buffer (10 mM Tris-HCl, pH 8.0, 200 mM NaCl, 1 mM EDTA and 0.5 mM EGTA) for 10 min. The lysis samples were resuspended and sonicated in sonication buffer (10 mM Tris-HCl, pH 8.0, 100 mM NaCl, 1 mM EDTA, 0.5 mM EGTA, 0.1% sodium deoxycholate and 0.5% N-lauroylsarcosine) using a Bioruptor Pico (Diagenode) with 30 s on and 30 s off for at least 6 cycles. After sonication, samples were centrifuged at 12,000 r.p.m. for 10 min at 4 °C, and 1% of sonicated cell extracts was saved as input. The resulting whole-cell extract was incubated with 5–10 µg of the appropriate antibody overnight at 4°C. After overnight incubation, antibody-bound whole-cell extracts were then incubated with Dynabeads Protein G (Invitrogen) for 1 h at 4°C and washed twice with sonication buffer, once with sonication buffer with 500 mM NaCl, once with LiCl wash buffer (10 mM Tris-HCl, pH 8.0, 1 mM EDTA, 250 mM LiCl and 1% NP-40) and once with TE with 50 mM NaCl. After washing, DNA was eluted in freshly prepared direct elution buffer (10 mM Tris-HCl (pH 8.0), 5 mM EDTA (pH 8.0), 300 mM NaCl and 0.5% SDS). Cross-links were reversed by overnight incubation at 65 °C. RNA and protein were digested using RNase A and proteinase K, respectively, and DNA was purified using UltraPure™ Phenol:Chloroform:Isoamyl Alcohol (Invitrogen) following the manufacturer’s instructions. For ChIP-qPCR assays, immunoprecipitated DNA was analyzed by quantitative real-time PCR and results were normalized relative to the amount of input DNA. The primer sequences for the qPCR reactions are listed in the key resource table. For ChIP-seq experiments, 100-300 bp DNA fragments were purified to prepare DNA libraries using NEBNext Ultra II DNA Library Prep Kit for Illumina following the manufacturer’s instructions. ChIP libraries were sequenced (50bp paired end reads) to at least 20 million reads/sample using an Illumina NextSeq 2000 or NovaSeq 6000 at the Genomic Resources Core Facility at Weill Cornell Medicine per manufacturer’s recommended protocol. The antibody used was from Cell Signaling Technology: STAT6 (5397S).

### CUT&RUN

CUT&RUN was performed as previously described ^39^ with minor adaptations. Cells were centrifuged for 3 min at 600 *g* at room temperature and then resuspended in wash buffer (20 mM HEPES pH 7.5, 150 mM NaCl, 0.5 mM Spermidine, 1x Roche Complete Protease Inhibitor EDTA-free tablet per 50 mL). Up to 500,000 cells per condition were centrifuged again for 3 min at 600 g at room temperature and resuspended in wash buffer. Activated Concanavalin-A beads (Bangs Laboratories BP531) in binding buffer (20 mM HEPES pH 7.5, 10 mM KCl, 1 mM CaCl2, 1 mM MnCl2) were added to each condition and incubated for 10 min on a rotator at room temperature. Beads with nuclei were kept on a magnetic stand and washed with Dig-wash buffer (0.05% digitonin in wash buffer). After discarding the liquid, beads were resuspended with indicated antibody in antibody buffer (2 mM EDTA in Dig-wash buffer) and incubated overnight at 4 °C on a nutator. Then beads were washed with freshly prepared Dig-wash buffer and incubated with 1 mg/ml pAG-MNase (CUTANA) in Dig-wash buffer for 1 h at 4 °C on a nutator. After two washes with Dig-wash buffer, ice-cold incubation buffer (2 mM CaCl2 in Dig-wash buffer) was added to the beads which were subsequently placed in a metal block in an ice-water bath maintained at 0 °C for 30 min. Freshly prepared 2x Stop buffer (340 mM NaCl, 20 mM EGTA, 4 mM EGTA, 0.05% digitonin, 100 µg/mL RNase A, 50 µg/mL glycogen) was then added and incubated for 10 min at 37 °C. Beads were placed on a magnet stand and supernatant was collected. 2 µL of 10% (wt/vol) SDS and 1.5 µL of 20 mg/ml Proteinase K were added to the supernatant and incubated for 30 min at 50 °C. DNA from the samples was purified using the Phase Lock Gel (Quantabio) and UltraPure™ Phenol:Chloroform:Isoamyl Alcohol (25:24:1, v/v) (Thermo Fisher Scientific). The antibodies for the reactions are listed in the key resource table. Sequencing libraries were prepared using NEBNext® Ultra II DNA Library Prep Kit. Libraries were sequenced (50bp paired end reads) to at least 10 million reads/sample on the Illumina Novaseq 6000 or Nextseq 500 at the Genomic Resources Core Facility at Weill Cornell Medicine. The antibodies used were from Bethyl Laborarotories: RelA (A301-824A), IRF1 (A700-039), from Millipore: trimethyl-Histone H3 (Lys27) (07-449), and from Abcam: Histone H3 (acetyl K27) (ab4729), Histone H3 (tri methyl K4) (ab8580).

### ATAC-seq, ChIP-seq and CUT&RUN analysis

Raw data underwent adapter trimming with FastP ^60^. Sequencing read alignments were performed against (GRCm38/mm10 for mouse and GRCh38/hg38 for human) reference genome using Bowtie2 ^61^. Peak calling was performed using MACS2 ^62^ with the following parameters: “--macs2 callpeak -f BAMPE nomodel shift −100 extsize 200 B SPMR -g hg38/mm10 -q 0.01.”. A reproducible ATAC-seq analysis pipeline (https://gitlab.com/hssgenomics/Shiny-ATAC) was used for differential peak analysis and ChromVAR analysis to evaluate motif enrichment ^63^. Footprint analyses were conducted using TOBIAS according to user manual ^64^. Motif databases in MEME format (JASPAR 2020 ^65^) were scanned against DNA sequences extracted from peak regions using bedtools ^66^. Significant motif matches were identified with a p-value threshold of 0.0001, and results were mapped back to the corresponding genomic regions for downstream analysis. Genomic tracks were visualized using Gviz ^67^.

### Motif enrichment analysis

*De novo* transcription factor motif analysis was performed with motif finder program *findMotifsGenome* from HOMER package ^68^ on given peaks. Peak sequences were compared to random genomic fragments of the same size and normalized G+C content to identify motifs enriched in the targeted sequences.

### Single-Cell RNA-Seq Data Collection and Processing

The single-cell RNA sequencing (scRNA-seq) data of two unwounded controls and three wounded samples were collected from Gene Expression Omnibus (GEO) under accession number GSE142471 ^32^. The Seurat package (v4) ^69^ was used to integrate all cells across samples. Cells that expressed less than 200 genes or more than 5000 were filtered. We further excluded cells that had > 12.5% mitochondrial gene expression. The highly variable genes were identified from these cells using Seurat with the default setting followed by principal component analysis (PCA). Dimensionality reduction across all datasets was achieved using Seurat’s *RunUMAP* function, employing an identical count of principal components (PCs) as in the clustering process for embedding calculation. Clusters were annotated using canonical cell type markers. Differentially expressed genes (DEGs) in cell types of clusters were identified by using the *FindMarkers* function embedded in Seurat package. In each comparison, genes with fold change > 1.5 and Benjamini-Hochberg adjusted p value <0.05 were considered statistically significant. To further identify subpopulations, we reanalyzed the hematopoietic cells separately, using the same workflow as described above. ClusterProfiler ^58^ was used to detect enriched pathways in the Reactome or Gene Ontology biological functions within each DEG set.

### Flow cytometric staining, sorting, and counting

Wounded skin samples were minced and digested with DNase I and collagenase II for 45 min in a rotator at 37 ℃. Cells were centrifuged to remove digestion buffer and red blood cells were lysed with ACK lysing buffer (Lonza) for 5 min. After washing with FACS buffer (PBS containing 0.5% BSA and 1 mM EDTA), cells were then filtered with 70 µm cell strainer and stained with LIVE/DEAD Fixable Blue Dead Cell Stain Kit (Thermo Fisher Scientific) in PBS. After washing with PBS, cells were stained with antibodies including F4/80 (BioLegend, BM8), Ly6G (BioLegend, 1A8), CD11b (BioLegend, M1/70), MHC II (BioLegend, M5/114.15.2), CD206 (BioLegend, C068C2) and CD45 (BioLegend, 30-F11). After washing, the cells were analyzed using BD FACS Symphony A3 Cell Analyzer and Flowjo software. Wounded skin cells for sorting were prepared following the method described above. The cells from 3 mice were pooled together and stained with antibodies listed above. Before sorting DAPI was added to the cells to label the dead cells. Cells were sorted using Becton-Dickinson Aria II. After sorting, the cells were quickly lysed with buffer RLT (QIAGEN).

### Adenovirus-mediated overexpression

Monocytes (1 × 10^6^ cells) were cultured in 24-well plates in complete RPMI-1640 medium supplemented with 10% heat-inactivated defined FBS, penicillin-streptomycin, L-glutamine, and 50 ng/mL human M-CSF for 5 d. The culture medium was replaced with fresh medium on the third day during the culture. On day 5 of culture, the old medium was removed and replaced with 200 μL of RPMI-1640 medium without penicillin-streptomycin with a final concentration of 2% heat-inactivated defined FBS and 50 ng/mL human M-CSF. Enhanced green fluorescent protein (GFP) or Ad-h-IRF1-HA, Ad-CMV-iKB (Vector Biosystems Inc) adenoviral particles (multiplicity of infection = 100 plaque-forming units/cell) were added to the cells accordingly. The cells were centrifuged at 1600 rpm for 30 min at room temperature (RT). After culturing for 12 h, 300 μL of complete RPMI-1640 medium supplemented with 10% heat-inactivated defined FBS, penicillin-streptomycin, L-glutamine, and 20 ng/mL human M-CSF was added to the cell culture. The cells transduced with Ad-h-IRF1-HA were cultured for an additional day before harvesting for experiments. The cells transduced with Ad-CMV-iKB were rested overnight prior to experiments.

### Statistical Analysis

Graphpad Prism for Windows or R (4.3.0) were used for statistical analysis. Information about the specific tests used, and number of independent experiments is provided in the figure legends. Two-way ANOVA with Sidak correction for multiple comparisons was used for grouped data. Otherwise, one-way ANOVA with the Geisser-Greenhouse correction and Tukey’s post hoc test for multiple comparisons was performed. The Mann-Whitney U test was utilized for comparison of sequencing data across groups. For paired data, paired t test was used. p-values of <0.05 were considered significant. ns: not significant, *p < 0.05, **p < 0.01, ***p < 0.001, ****p < 0.0001.

## ACKNOWLEDGMENTS

We would like to thank previous and current members of the Ivashkiv Laboratory. We also thank the Genomics Resources Core Facility at Weill Cornell Medicine for next generation sequencing, the Weill Cornell Medicine – HSS Flow Cytometry Core Facility for flow cytometry support, and Yurii Chinenov (HSS Genomics Center) for advice and discussions.

## Author contributions

R.Y. conceptualized, designed, and performed most of the experiments, performed bioinformatic analysis, prepared figures and wrote the manuscript. B. M., C. Y. and R. B. contributed experiments or experimental expertise; D.O. performed bioinformatic analysis. L.B.I. conceptualized and oversaw the study and edited the manuscript. All authors reviewed and provided input on the manuscript.

## Data and materials availability

### Lead contact

Further information and requests for resources and reagents should be directed to and will be fulfilled by the Lead Contact, Lionel B. Ivashkiv.

### Materials availability

Materials from this study will be available from the lead contact upon request.

### Data and code availability

- Sequencing data from this study have been deposited at GEO (GSE252738 and GSE287605) and are publicly available from the date of publication, reviewers can access it with secure token: kjstqswcbrgpncz (GSE252738) and unqfcggwfxsxlsx(GSE287605).
- Original code has been deposited at gitlab and is publicly available from the date of publication. Urls: https://gitlab.com/hssgenomics/Shiny-ATAC.
- Any additional information required to reanalyze the data reported in this paper is available from the lead contact or first author on request.

