## Supplementary figures for "Beyond Antagonism: IL-4 Exploits TNF signaling to Shape Its Gene Expression Signature in Monocytes and Macrophages"

Yuan et al., Extended Data Fig 1

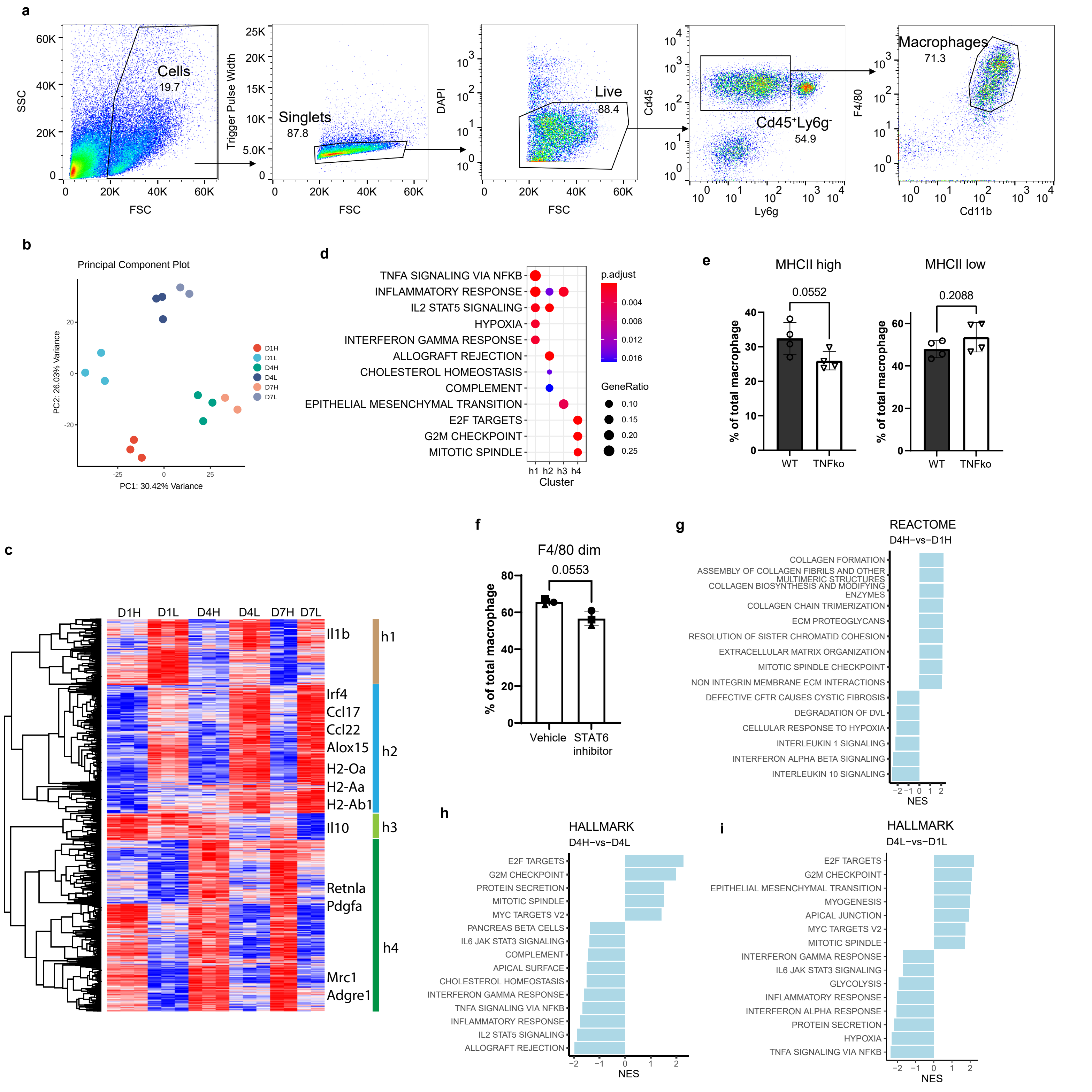

Yuan et al., Extended Data Fig 2

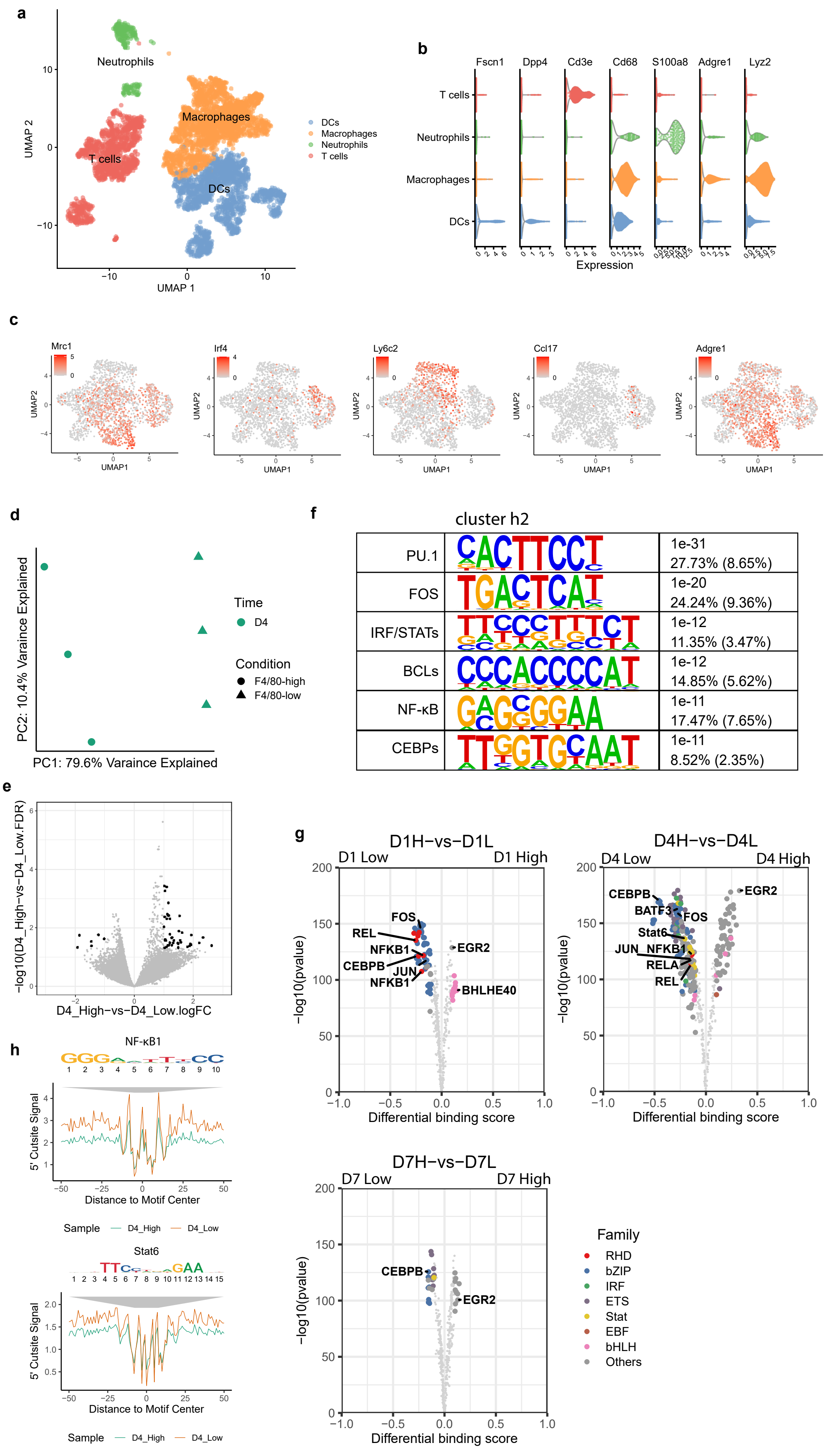

**Yuan et al., Extended Data Fig. 3**

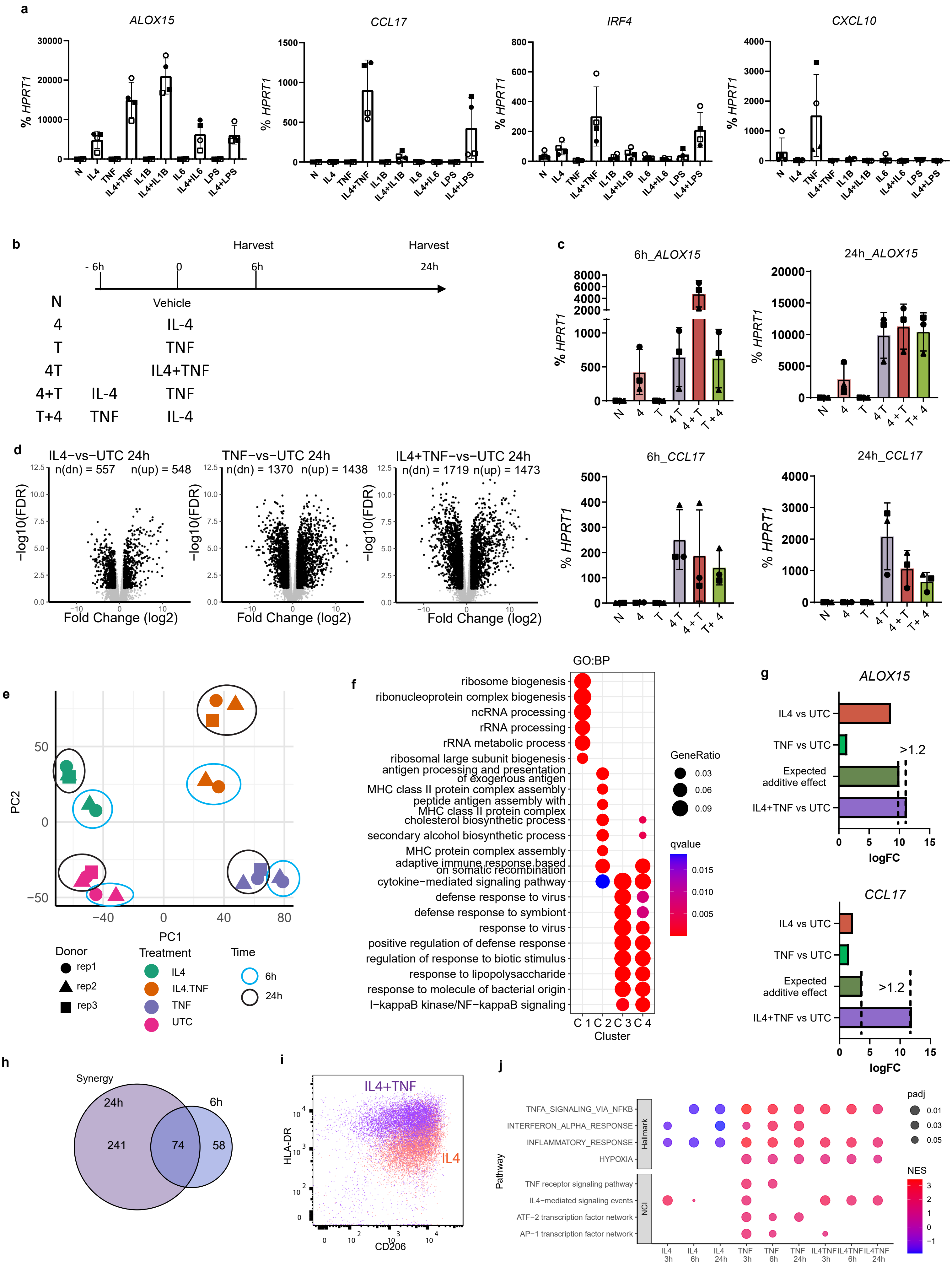

**Yuan et al., Extended Data Fig. 4**

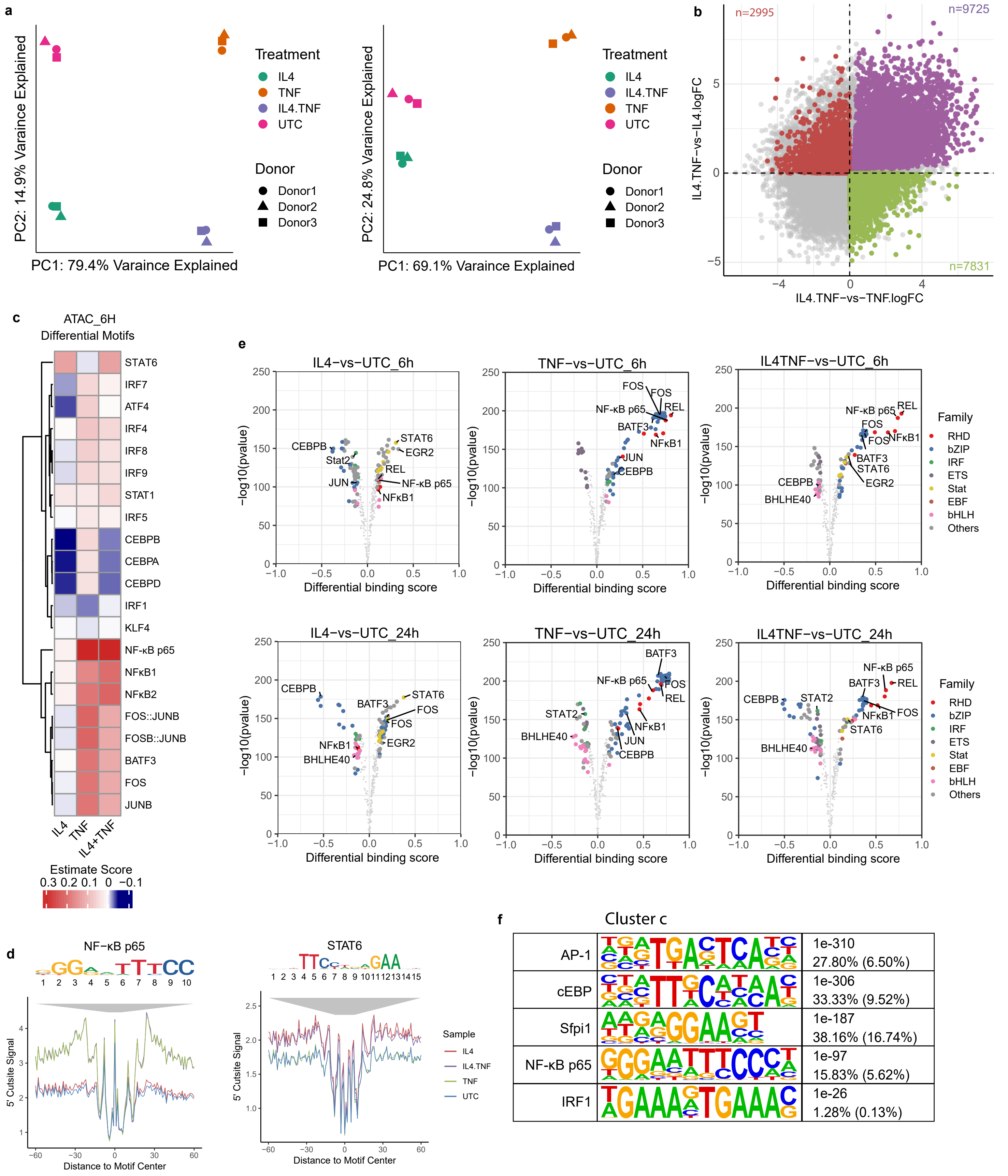

Yuan et al., Extended Data Fig. 5

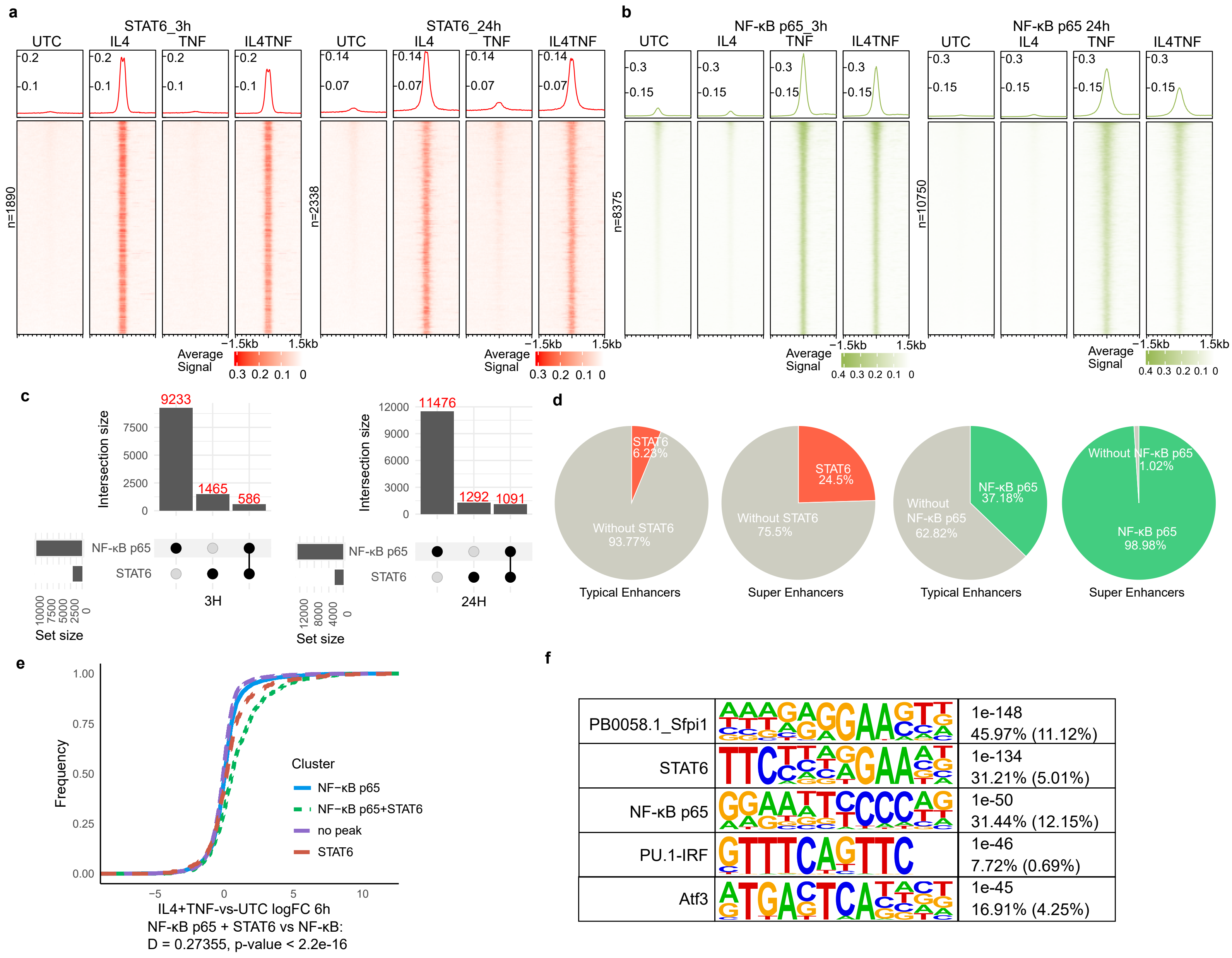

**Yuan et al., Extended Data Fig. 6**

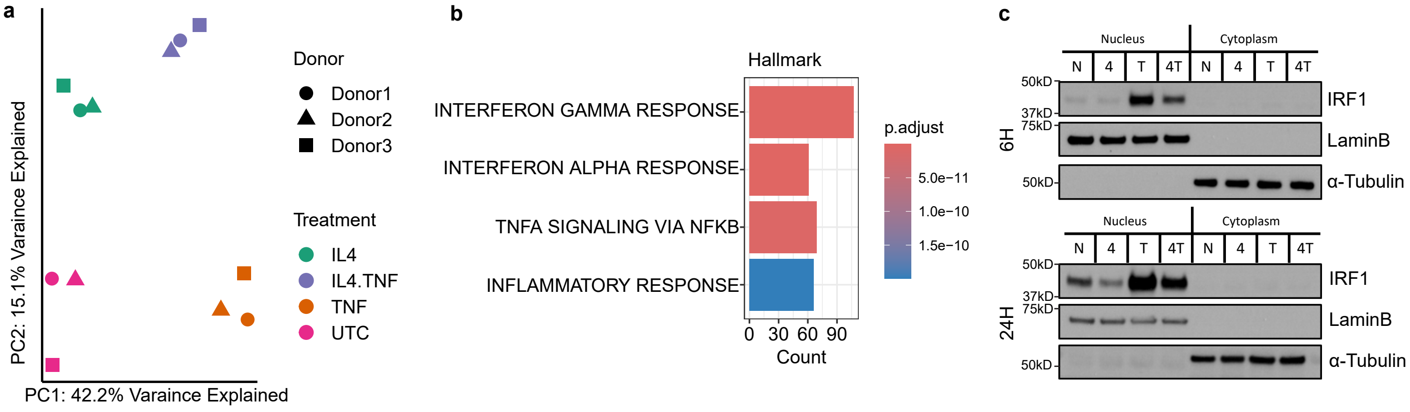
