## Supplementary figure legends for "Beyond Antagonism: IL-4 Exploits TNF signaling to Shape Its Gene Expression Signature in Monocytes and Macrophages"

**Extended Data Figures Legends**

**Extended Data Fig 1. Expression of (TNF + IL-4)-induced genes in skin wound macrophages.**

**a**, Flow cytometry gating strategy for detection of macrophages in mouse skin wounds at 4 dpi.

**b**, Principal component analysis (PCA) of top 2000 variable genes from RNA-seq data (D1: n=3; D4: n=3; D7: n=2).

**c**, Hierarchical clustering was performed using Euclidean distance on 1937 differentially up-regulated genes (log2 fold change > 1 and FDR < 0.05) in any pairwise comparison between F4/80-high and F4/80-low macrophages. The dendrogram was analyzed to determine an appropriate cut point, resulting in four clusters. Values from biological replicates are shown in separate columns for each sample (D1: n=3; D4: n=3; D7: n=2).

**d**, Hallmark pathway enrichment analysis of the gene clusters in **c**.

**e**, Flow cytometric analysis of MHCII high and MHCII lo macrophages isolated from skin wounds at 4 dpi (n=4). Results are cumulative from two independent experiments. Paired t test was used. Data presented as means ± SD.

**f**, Flow cytometric analysis of F4/80 dim macrophage population isolated from skin wounds at 4 dpi (n=3) treated with vehicle or STAT6 inhibitor. Paired t test was used. Data presented as means ± SD.

**g-i**, GSEA analysis of gene lists, ranked by log2 fold change (log2FC), under specified comparisons and time points (**g**, D4H vs D1H; **h**, D4H vs D4L; **i**, D4L vs D1L). Analysis includes Reactome (**g**) and Hallmark (**h**, **i**) gene sets.

**Extended Data Fig 2. scRNA-seq and ATAC-seq reveals distinct macrophage clusters.**

**a**, UMAP plot showing unsupervised clustering of scRNA-seq data obtained from GSE142471 of cells from skin wounds at 4 dpi. Plot shows the UMAP projection of integrated identification of cell types based on canonical markers.

**b**, Violin plots of representative hematopoietic cell markers across the cell clusters identified in 2c.

**c**, UMAP plots showing expression of indicated genes using log2-transformed normalized counts.

**d**, PCA of top 2000 variable peaks from RNA-seq data (D4: n=3).

**e**, Volcano plots display the statistical significance (-log10 FDR, y-axis) versus the log2 fold change of ATAC-seq data, highlighting selected peaks that with more signal strength in F4/80-high macrophages (right) and F4/80-low macrophages (left) at 4 dpi.

**f**, The most significantly enriched transcription factor (TF) motifs identified by *de novo* motif analysis using HOMER in enhancer peaks associated with genes in cluster h2 from Extended Fig. 1c.

**g**, Volcano plots showing differentially occupied motifs identified by TOBIAS, colored by different transcription factor binding motif families at 1, 4 and 7 dpi. Right: motifs occupied in F4/80-high macrophages. Left: motifs occupied in F4/80-low macrophages.

**h**, The signal enrichment profiles of NF-κB and Stat6 motifs at 4 dpi using ChromVAR.

**Extended Data Fig 3. Synergistic and Antagonistic Interplay between IL-4 and TNF-α in Regulating Gene Expression in Human Monocytes.**

**a**, RT-qPCR analysis of expression of the indicated genes in human $CD14^+$ monocytes co-stimulated with IL-4 and various pro-inflammatory mediators (TNF: 10ng/mL, IL-1β: 10ng/mL, LPS: 10ng/mL) for 24 hours (n=4). Error bars represent SD.

**b**, Experimental design: human CD14$^+$ monocytes were stimulated with IL-4 (4), TNF (T), co-stimulation of IL-4 and TNF (4T), pretreated with IL-4 for 6 hours (4+T) or pretreated with TNF for 6 hours (T+4). Monocytes were treated for 6 hr or 24 hr.

**c**, RT-qPCR analysis of expression of the indicated genes in human CD14$^+$ monocytes stimulated with indicated treatments and times as shown in 1**b** (n=3). Error bars represent SD.

**d**, Volcano plots of statistical significance (-log10 FDR) plotted against log2 ratio differential gene expression in RNA-seq data showing differentially induced (right) or suppressed (left) genes after individual or combined treatment with IL4 and TNF compared to untreated control (UTC) monocytes. Significantly DEGs (log2 fold change > 1; FDR < 0.05) (black), non-significant genes (grey).

**e**, PCA of top 1000 variable genes from RNA-seq data (6h: n=2; 24h: n=3).

**f**, Enriched Gene Ontology (GO) Biological Process (BP) categories of the genes in Fig. 1c.

**g**, Bar plot depicting the definition of synergy genes based on a predefined cutoff. Synergy is defined as > 1.2-fold relative to the expected additive effect defined by addition of (IL-4 vs UTC) and (TNF vs UTC).

**h**, Venn diagram of the overlap between synergy genes and "*de novo*" induced genes at 24 hours. The "*de novo*" genes exhibit a log fold change (logFC) of less than 1 after individual IL-4 and TNF treatments at 24 hours compared to untreated control, but a logFC greater than 1 with a false discovery rate (FDR) below 0.05 after IL-4 and TNF co-stimulation.

**i**, Representative flow cytometry plot of human monocyte subsets defined by MHC II and CD206 cell surface expression after 24 hr stimulation. (IL-4: red, IL-4+TNF: purple). Cells were gated on live singlets (using a FSC-W versus FSC-A gate).

**j**, GSEA analysis of gene lists, ranked by log2 fold change (log2FC), under specified treatments and time points, compared to untreated controls. Analysis includes known Hallmark gene sets and gene sets from the Nature Pathway Interaction Database (NCI).

**Extended Data Fig 4. Regulation of chromatin accessibility and transcription factor activity.**

**a**-**f**. Analysis of the same ATAC-seq data as Fig. 3, n = 3 at each time point.

**a**, Principal component analysis (PCA) of top 1000 variable ATAC-peaks at 6 hours (left panel) or 24 hours (right panel).

**b**, Dot plot showing ATAC peaks differentially induced (FDR < 0.05) by co-stimulation relative to IL-4 or TNF alone. Purple, peaks significantly up-regulated only by co-stimulation. Red, upregulated peaks unique to IL-4 stimulation; green, upregulated peaks unique to TNF stimulation; grey: non-significant peaks.

**c**, Heatmap of differentially enriched motifs at 6 hours comparing cytokine treatments to untreated controls from signal enrichment around specific motif analysis using ChromVAR.

**d**, Enriched motif profiles of NF-κB p65 and STAT6 at 6 hours by ChromVAR.

**e**, Volcano plots showing differentially induced (right) or suppressed (left) motifs identified by TOBIAS, colored by different motif families. Left panel: 6 hours, right panel: 24 hours.

**f**, *De novo* motif analysis using HOMER of peaks in Cluster c in Fig. 3e.

**Extended Data Fig 5. Cooperation between NF-κB and STAT6 to enhance gene responses**.

**a**-**f**. Analysis of same ATAC-seq data set as in Fig. 4, n = 3.

**a**, **b**, CUT&RUN and ChIP-seq analysis of NF-κB and STAT6 in human monocytes treated with indicated cytokines for 3 or 24 hours. Data are presented as normalized signal density ± 2.5kb around peak centers. Color scale represents the depth-normalized counts of peaks.

**c**, UpSet Plot of STAT6 and NF-κB p65 overlapping peaks at 3 and 24 hours. Intersection and unique occurrences of binding peaks of STAT6 and NF-κB p65 transcription factors. The plot's matrix layout presents the overlap and exclusivity of these peaks. Vertical bars indicate the number of peaks at each time point, horizontal bars represent the total number of peaks identified for each factor.

**d**, Pie charts showing the percentage of STAT6 and NF-κB p65 binding sites identified as occurring in typical enhancers or super enhancers based on H3K27Ac signal strength calculated in Fig. 4d.

**e**, ECDF plot with a Kolmogorov-Smirnov statistical test at 6 hours. The x-axis displays the log2 fold change (log2FC) of gene expression, representing the magnitude and direction of gene expression changes under different conditions. The y-axis quantifies the cumulative frequency of genes at each log2FC level. Four categories of genes: genes with only NF-κB p65 peaks, with only STAT6 peaks, with NF-κB-p65 and STAT6 co-binding, and with no binding of these two transcription factors. The D value of Kolmogorov-Smirnov test indicates the significance of differences of the log2FC between selected comparison (genes with NF-κB-p65 and STAT6 co-binding versus genes with only NF-κB p65 peaks). Values of $p < 0.05$ were considered statistically significant.

**f**, *De novo* motif analysis results using HOMER of STAT6 and NF-κB p65 co-binding sites.

**Extended Data Fig 6. IL-4 targets IRF1-binding genomic elements to suppresses TNF-induced interferon responses**.

**a**, PCA plot of top 1000 variable IRF1 peaks at 24 hours after cytokine stimulation.

**b**, Gene set over-representation analysis of the genes associated with IRF1 binding sites using Hallmark gene sets. Values of $q < 0.05$ were considered statistically significant.

**c**, Immunoblot of IRF1, Lamin B1 or α-Tubulin using nuclear extracts or cytosolic extracts at 6 hours or 24 hours after indicated stimulation. One representative experiment out of 3 (6 hours) or 2 (24 hours) is shown.
